# Control of pancreatic islet function and glucose homeostasis by a novel microexon program misregulated in type 2 diabetes

**DOI:** 10.1101/2022.04.02.486809

**Authors:** Jonàs Juan-Mateu, Simon Bajew, Marta Miret-Cuesta, Luis P. Íñiguez, Amaya López-Pascual, Sophie Bonnal, Goutham Atla, Sílvia Bonàs-Guarch, Jorge Ferrer, Juan Valcárcel, Manuel Irimia

## Abstract

Pancreatic islets control glucose homeostasis by the balanced secretion of insulin and other hormones, and their abnormal function causes diabetes or hypoglycemia. Here, we uncover a conserved program of alternative microexons included in mRNAs of islet cells, particularly in genes involved in vesicle transport and exocytosis. Islet microexons (IsletMICs) are regulated by the RNA binding protein *SRRM3* and represent a subset of the larger neural program that are particularly sensitive to the levels of this regulator. Both *SRRM3* and IsletMICs are induced by elevated glucose levels, and depletion of *SRRM3* in beta cell lines and mouse islets, or repression of particular IsletMICs using antisense oligonucleotides, leads to inappropriate insulin secretion. Consistently, *SRRM3* mutant mice display defects in islet cell identity and function, leading to hyperinsulinemic hypoglycemia. Importantly, human genetic variants that influence *SRRM3* expression and IsletMIC inclusion in islets are associated with fasting glucose variation and type 2 diabetes risk.

## Introduction

Pancreatic islets are complex endocrine micro-organs that play an essential role in maintaining glucose homeostasis through the balanced secretion of hormones into the bloodstream. Insulin, the major glucose-lowering hormone, is produced exclusively by pancreatic islet beta cells. Alterations in beta cell function and insulin secretion perturb glucose homeostasis, leading to either diabetes, when there is a deficiency of insulin, or to persistent hypoglycemia, when insulin is secreted in excess. Over the last decade, extensive research has uncovered cell type-specific transcriptional programs driving islet cell differentiation and identity^1^ as well as transcription factors and signaling pathways involved in the acquisition of functional maturity and the ability to regulate insulin secretion^2^. Moreover, integration of transcriptomic, epigenomic and genetic data has shown that dysregulation of islet-specific gene expression is a common mechanism underlying type 2 diabetes (T2D)^3, 4^ and monogenic forms of diabetes^5^.

In contrast, the roles of alternative splicing in pancreatic islet function remain not well understood. Alternative splicing regulates the differential processing of introns and exons to generate multiple mRNA isoforms from a single primary transcript, contributing to transcriptomic and proteomic diversity. It is estimated that over 95% of human multi-exonic genes undergo alternative splicing^6, 7^, a substantial fraction of which is regulated in a tissue-and cell-type enriched manner^8^. A plethora of studies have provided experimental evidence that alternative splicing regulation is essential for many biological processes, and unveiled the roles of several RNA binding splicing regulatory proteins in tissue-specific development and functions^9^. In the case of pancreatic beta cells, previous findings indicate that they express neural-related RNA binding proteins that impact the regulation of insulin secretion^10, 11^, and that splicing misregulation contributes to beta cell apoptosis in an *in vitro* model of type 1 diabetes^12^. However, the presence, regulation and impact of islet-enriched alternative splicing programs on endocrine cell development and identity remain unexplored.

Microexons, very short exons (between 3 to 27 nucleotides), constitute a unique form of tissue-enriched alternative splicing, and were originally described as a highly conserved program preferentially included in neurons^13, 14^. These microexons tend to maintain the open reading frame, introducing as few as one amino acid into the surface of proteins, where they often rewire protein-protein interactions^13^. Microexon inclusion in neurons is largely dependent on the RNA binding proteins SRRM3 and SRRM4, which provide essential functions in nervous system development^15–18^. Both factors share a common 39 amino acid domain (enhancer of microexons, or eMIC) at their C-termini that is necessary and sufficient for microexon inclusion^19^. Interestingly, microexons are misregulated in the brains of some autistic patients^13, 20^.

Here, we found that a subset of neural microexons is also present in pancreatic islets, forming an evolutionarily conserved program that particularly affects genes involved in insulin secretory function and T2D risk. These microexons respond to physiological changes in glucose levels and are regulated by *SRRM3*. Consistently, misregulation of microexons in beta cell lines, either globally through *SRRM3* depletion or individually using antisense oligonucleotides, result in dysregulated insulin release. In line with this, *Srrm3* knockout mice show alterations in islet cell identity, increased insulin secretion and persistent hypoglycemia. Furthermore, genetic variations affecting the regulation of *SRRM3* and microexon inclusion in human islets are associated with fasting glucose and T2D risk. Our results thus uncover a post-transcriptional regulatory program necessary for mature islet cell function and glycemic control.

## Results

### Human and rodent islets express a subset of neural microexons

To systematically identify tissue-enriched alternative exons in human islets, we profiled alternative splicing genome-wide using publicly available RNA-seq data from islets, endocrine pancreatic cells and multiple neural and non-neural tissues, including exocrine pancreas (Supplementary Table 1), and searched for exons enriched in endocrine pancreatic and/or neural tissues compared to other tissues (see Methods). Surprisingly, out of 201 exons enriched in endocrine pancreas, 109 (54.2%) consisted of 3-27 nucleotide microexons (hereafter IsletMICs) (Fig. 1A and Supplementary Table 2). Interestingly, endocrine pancreas displayed the highest ratio of microexons to longer tissue-specific exons found in any human tissue (e.g. compared to 33% in neural tissues, 11% in skeletal muscle, 6% in liver or 1% in exocrine pancreas). The vast majority (106/109, 97.3%) of IsletMICs were also highly included in neural cells, and thus represent a subprogram that is nested within the larger neural microexon program (Fig. 1A,B). IsletMICs were highly included in the different types of endocrine cells, as observed using both bulk (Fig. 1C) and single cell RNA-seq data (Extended Data Fig. 1A,B), but were largely absent or lowly included in other non-neural tissues, including exocrine pancreas acinar and ductal cells (Fig. 1A,C and Extended Data Fig. 1A,B). As is the case for neural microexons^13^, IsletMICs showed a high degree of conservation from human to rodents at both the genomic and regulatory level (Fig. 1D), and largely maintained the open reading frame and are thus predicted to generate alternative protein isoforms (Extended Data Fig. 1C). These patterns were in sharp contrast with those of the program of longer islet exons (hereafter IsletLONGs), which showed a lower overlap with neural-enriched exons (Extended Data Fig. 1D,E), lower evolutionary conservation (Fig. 1D) and a higher frequency of open reading frame-disrupting events (Extended Data Fig. 1C).

**Figure 1.**
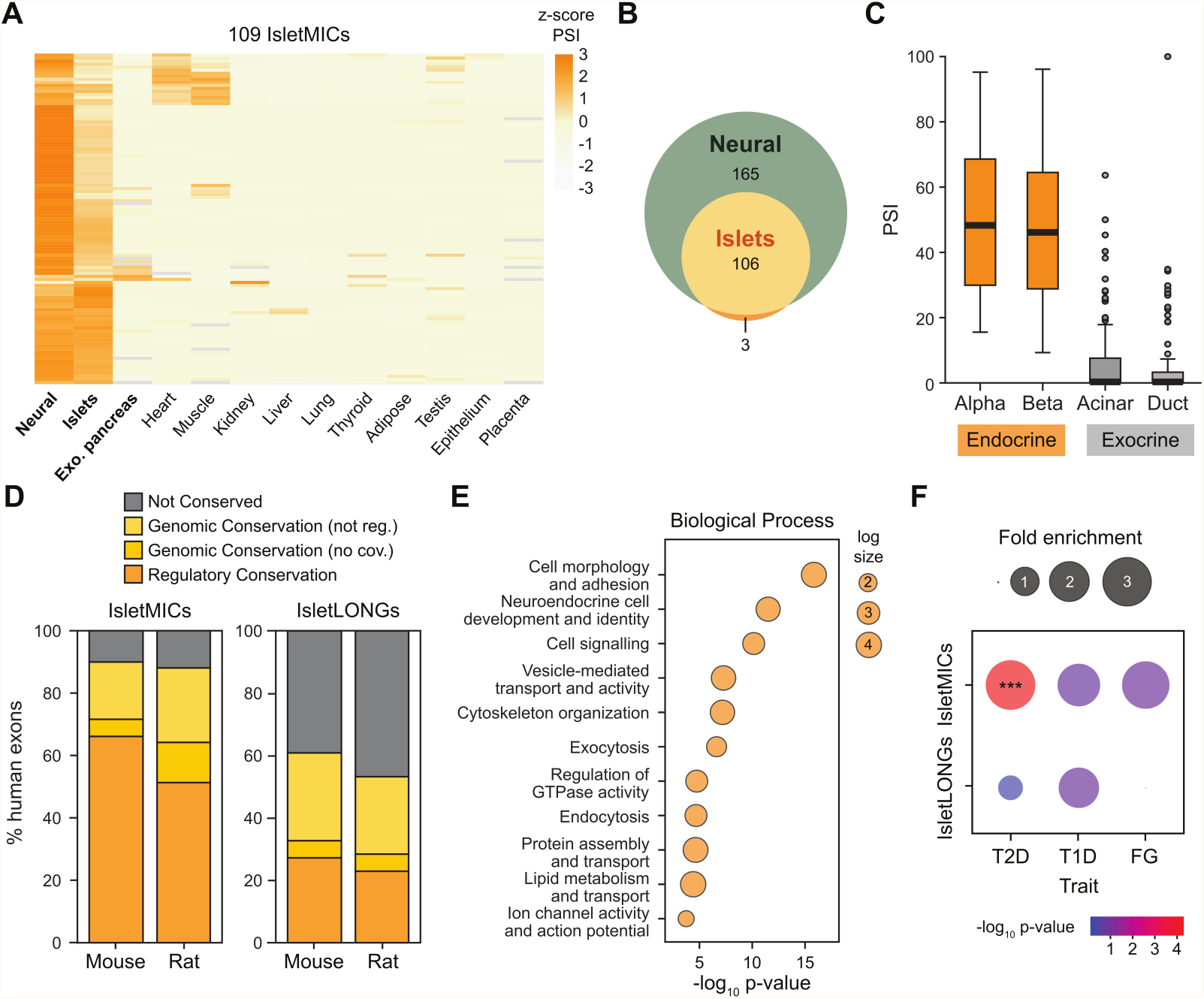
Pancreatic islets express a conserved subset of neural microexons. **A)** Heatmap of tissue inclusion levels (expressed as z-score of the percent-spliced-in [PSI]) of a subset of human microexons (≤27 nt) enriched in pancreatic islets (IsletMICs). PSI values are the mean of three to ten tissue samples per group. **B)** Overlap between IsletMICs and neural-enriched microexons illustrates how the IsletMICs program is nested within the larger neural program. **C)** Mean inclusion levels of IsletMICs in different cell types from endocrine (beta and alpha cells, orange) and exocrine (acinar and duct cells, grey) pancreas from bulk RNA-seq. PSI values are the mean of four samples per group. **D)** Evolutionary conservation of human IsletMICs at the genomic and tissue-regulatory level compared with mouse and rat. Regulatory conservation refers to exons with enriched inclusion in endocrine pancreas compared to other tissues in the target species. Genomic conservation refers to exons having an ortholog in the target species, but either no specific islet enrichment (“not reg.”) or not sufficient read coverage to assess regulation (“no cov.”). **E)** Top ten enriched GO terms for human IsletMIC-harboring genes. P-values were corrected for FDR. **F)** Enrichment of islet-enriched exons in genes harboring T2D, T1D or fasting glucose (FG) risk variants. The size of the circle is proportional to ratio of the observed versus expected overlaps. P-values correspond to hypergeometric tests (color coded). ***, p < 0.001 versus all multi-exonic genes with matched expression.

Genes hosting IsletMICs were enriched in a wide array of processes associated with endocrine secretory functions (Fig. 1E), including regulation of GTPase activity (*ACAP2, CPEB2, GIT1, RAPGEF6*), ion channel activity (*CACNA2D1, CACNA1D*), exocytosis (*CASK, SYNJ1, PPFIA3*) and vesicle-mediated transport (*VAV2, KIF1B, DCTN1*). Remarkably, in contrast to IsletLONGs and to NeuralMICs, IsletMICs were significantly enriched in genes located in T2D susceptibility loci (Fig. 1F; e.g. *TCF7L2, PTPRS, MYH10, CPEB3, VTI1A*). Together, these data point to potentially important roles of the IsletMIC program in pancreatic islet development and/or function.

### IsletMICs are regulated by the splicing factor *SRRM3*

To investigate the regulation of IsletMICs, we examined mRNA expression of the well-established microexon regulators *SRRM4* and *SRRM3* in pancreatic islets. Whereas human and rodent islets showed little or undetectable expression of the master neuronal microexon regulator *SRRM4* (Figs. 2A and Extended Data Fig. 2A,B), mRNA expression of its paralog *SRRM3* was readily detectable in human, mouse and rat islets, although at consistently lower levels than in neural tissues (Figs. 2A and Extended Data Fig. 2A,B). To test whether *SRRM3* regulates the inclusion of IsletMICs, we performed siRNA-mediated knock-down (KD) of *SRRM3* transcripts in human EndoC-βH1 and rat INS-1E beta cell lines, followed by analysis of the inclusion of selected microexons by RT-PCR as well as global RNA-seq analyses. A ∼50% reduction of *SRRM3* (Figs. 2B and Extended Data Fig. 2C) was sufficient to induce skipping of most IsletMICs (ΔPSI ≤ −15) in both human and rat beta cells (Figs. 2C-E and Extended Data Fig. 2D-F, Supplementary Table 3). Moreover, short exons comprised the majority of misregulated alternatively spliced exons upon *Srrm3* KD (Fig. 2D and Extended Data Fig. 2E), supporting a highly specific function for *SRRM3* in the inclusion of IsletMICs.

**Figure 2.**
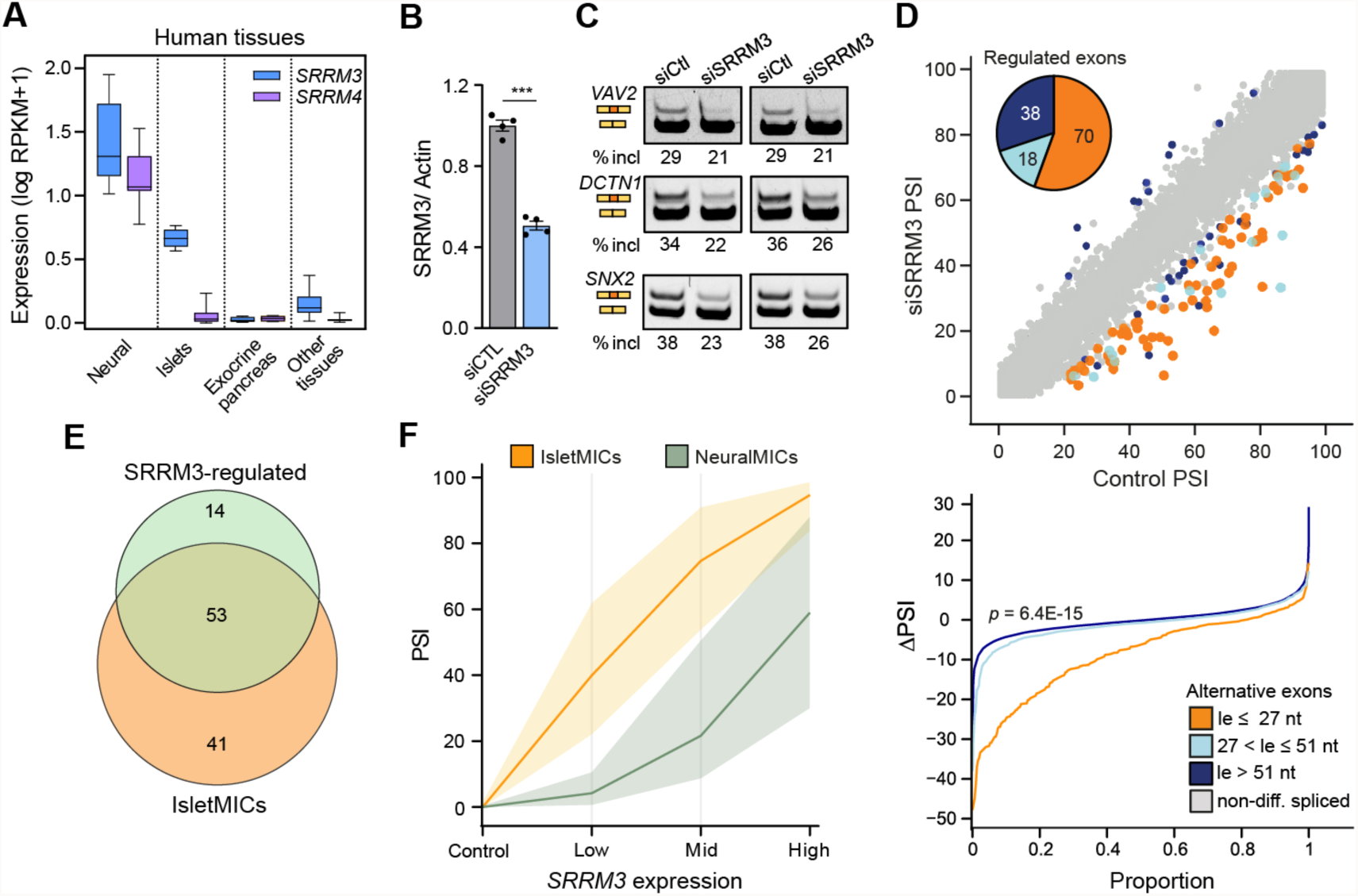
IsletMICs correspond to the subset of neuronal microexons with high sensitivity to *SRRM3*. **A)** mRNA expression of the neural microexon regulator *SRRM4* and of its paralog *SRRM3* in neural tissues, pancreatic islets, exocrine pancreas and other human tissues (data from VastDB, *n* = 4-12 tissue samples per group). **B-D)** siRNA-mediated knockdown (KD) of *SRRM3* in EndoC-βH1 human beta cell line. **B)** *SRRM3* mRNA levels measured by qPCR following 48h transfection with Srrm3 (siSRRM3) or control (siCTL) siRNAs, normalized to actin (*n* = 5, mean ± s.e.m). ***, p < 0.001 versus siCTL; paired t test. **C)** RT-PCR assays of selected microexons in control and *SRRM3* KD cells. The positions of inclusion/skipping isoforms and the percent of microexon inclusion are indicated for two biological replicates. **D)** Global impact of *SRRM3* KD on exon inclusion levels estimated by PSI values from RNA-seq data. Differentially included exons in *SRRM3* KD vs control are shown in orange (microexons, length [le] ≤ 27 nt), light blue (short exons, length between 28 and 51 nt) and dark blue (cassette exons, length > 51 nt). The pie chart shows the number of misregulated exons (|ΔPSI| ≥ 15) according to their size range. Lower panel shows ΔPSI cumulative proportions for microexons, short exons and alternative cassette exons. P-value was obtained from two-sided Wilcoxon test comparing the distributions of microexons and cassette exons of length > 51 nt. PSI values are the mean of three independent experiments. **E)** Overlap between *SRRM3*-regulated microexons in human EndoC-βH1 cells (*SRRM3*-regulated) and human IsletMICs. Only microexons with sufficient read coverage in both comparisons are shown. **F)** *SRRM3* overexpression at three different levels in HeLa cells reveals different sensitivities to *SRRM3*-mediated regulation for IsletMICs and neuronal-only MICs (NeuralMICs). Mean inclusion levels for IsletMICs and NeuralMICs in control cells and at three different *SRRM3* expression levels are shown (*n* = 1). Area fill represents the interquartile range.

Because IsletMICs mainly comprised a defined subset of the neural microexon program (Fig. 1B), we next wondered how this two-level (neural-islet) tissue-enriched regulation was achieved. Given that the combined levels of *SRRM3/4* were significantly lower in islets than in neural tissues (Fig. 2A and Extended Data Fig. 2A,B), we hypothesized that the reason why IsletMICs are included in both tissues is because they are more sensitive to low levels of *SRRM3/4* expression than neural-only microexons, whose inclusion would require higher levels of the regulators. To test this hypothesis, we stably expressed *SRRM3* in HeLa cells, a non-neural cell line in which microexons are not included, at three different levels (low, mid and high) using a doxycycline-inducible system. As predicted, whereas most IsletMICs were highly included already at low levels of *SRRM3* expression, the majority of neural-only microexons required higher *SRRM3* expression to reach detectable inclusion levels (Fig. 2F and Supplementary Table 3). Comparing the changes in microexon inclusion between IsletMICs and neural-only microexons at each induction level further evidenced that IsletMICs displayed higher responsiveness to SRRM3 (Extended Data Fig. 2H). These findings strongly argue that differential intrinsic sensitivity of microexons to SRRM3/4 activity is the major determinant of their tissue-enriched inclusion pattern, and that the two-level expression of *SRRM3/4* in islets and neural tissues is sufficient to configure two nested tissue-enriched microexon programs.

### *Srrm3* depletion affects insulin secretory functions of beta cells at multiple levels

To gain insights into the functional role of IsletMICs in pancreatic beta cells, we next assessed the impact of *SRRM3* depletion on insulin secretion. Remarkably, *SRRM3* KD led to a significant increase in insulin release following static stimulation with high glucose, with or without the adenylyl cyclase activator forskolin, in human Endoc-βH1 and rat INS-1E beta cells (Fig. 3A,B). Dynamic insulin secretion measurements in INS-1E cells showed a significant increase in both the first (0-15 min) and the second (16-40 min) phases of insulin release (Fig. 3C), suggesting that *Srrm3* silencing impacts both the K-ATP channel-dependent pathway responsible for the first phase, as well as other K-ATP-independent pathways contributing to the second phase^21^. In line with this, exposure of cells to the ATP-sensitive K+ channel inhibitor tolbutamide, in the presence of low or high glucose, revealed that *Srrm3* KD increased insulin release in both conditions (Fig. 3D).

**Figure 3.**
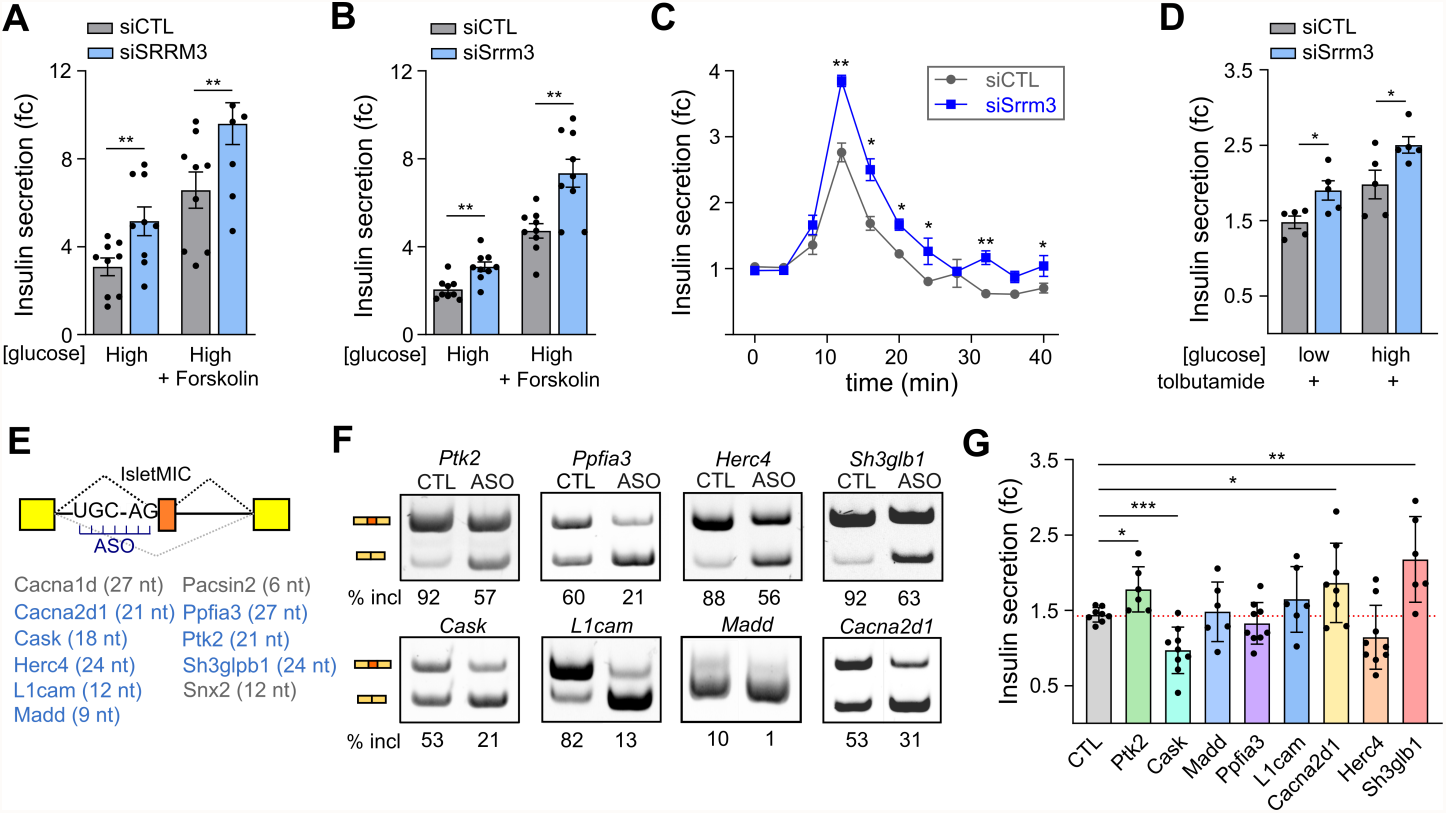
*Srrm3* regulates insulin secretory functions of pancreatic beta cells. Human EndoC-βH1 (**A**) or rat INS-1E (**B-D**) beta cells were transfected with control (siCTL) or *SRRM3*-targeting (siSRRM3) siRNAs for 48h. Increased basal and stimulated secretion upon *SRRM3* KD after exposure to low and high glucose concentrations with or without the cAMP activator forskolin in EndoC-βH1 (A) and INS-1E cells (B) (*n* = 9). **C)** Increased dynamic insulin secretion upon Srrm3 KD; measurements were made every 4 min after replacing 1.7 mM glucose medium with 17 mM glucose, (*n* = 3). **D)** Static insulin secretion after exposure to the K-ATP channel inhibitor tolbutamide, in combination with low and high glucose, (*n* = 5). **E)** Scheme indicating the design of ASOs complementary to pre-mRNA sequences encompassing the UGC *Srrm3*-responsive motif and the 3’ss. Targeted genes and microexon sizes are shown. Grey indicates cases were no significant changes in PSI were achieved. **F)** RT-PCR analysis of microexon inclusion/exclusion after transfection with control (CTL) or microexon-specific (ASO) ASOs in rat INS-1E cells. **G)** Glucose-stimulated insulin secretion following transfection with microexon-specific ASOs. Glucose stimulation index indicates the fold change insulin secretion between 1.7 mM and 17 mM glucose conditions (*n* = 6-9). All data are presented as mean ± s.e.m. Statistical comparisons: *, p < 0.05; **, p < 0.01 and ***, p < 0.001 versus siCTL or CTL ASO; unpaired t test.

Because genes with IsletMICs were enriched in multiple functions that may impact insulin secretion (Fig. 1E), we next investigated different relevant cellular pathways involved in this process upon *Srrm3* depletion. First, to test whether alterations in cell metabolism contributed to the observed increase in insulin secretion, we transfected INS-1E cells with a cytoplasmic ATP FRET probe ^22^ and measured ATP levels at resting (1.7 mM glucose) and stimulatory (17 mM glucose) conditions in *Srrm3*-depleted and control cells (Extended Data Fig. 3A). *Srrm3*-depleted cells showed a two-fold increase in ATP levels in low glucose conditions (Extended Data Fig. 3A–B), and faster ATP production following stimulation with high glucose (Extended Data Fig. 3C), suggesting an impact of *Srrm3* on cellular energy metabolism. Second, we evaluated alterations on the organization and remodeling of cytoskeleton, which plays crucial roles in insulin granules exocytosis^23^. *Srrm3*-depleted INS-1E cells showed a visible change in cell morphology, and staining of filamentous actin revealed a significant reduction of actin-rich filopodia (Extended Data Fig. 3D–F). Moreover, the actin cytoskeleton showed altered remodeling dynamics when stimulated with high glucose as measured by the intensity ratio between filamentous and globular actin (Extended Data Fig. 3G). Along similar lines, *Srrm3*-depleted cells under low glucose displayed decreased tubulin staining, consistent with alterations in microtubule polymerization (Extended Data Fig. 3H,I). Collectively, these observations argue that *Srrm3* impacts multiple aspects of the process of insulin secretion.

Given this diversity of phenotypes, we next tested whether misregulation of individual IsletMICs was sufficient to induce changes in glucose-stimulated insulin secretion. To that end, we designed antisense oligonucleotides (ASO) harboring 2′-O-methyl phosphorothioate modifications targeting microexon 3′ splice sites and adjacent regulatory sequences, including the UGC motif required for SRRM3-mediated regulation^18, 24^ (Fig. 3E). We selected 11 IsletMICs dependent on *SRRM3* in both human and rat cell lines, which were located in genes belonging to different pathways previously linked to beta cell function, membrane depolarization and/or exocytosis. In 8 out of 11 cases, we were able to induce microexon skipping to an extent similar to that observed following *Srrm3* KD (Fig. 3F). Importantly, microexon skipping led to a significant misregulation of glucose-stimulated insulin secretion in four cases (Fig. 3G), three of which enhanced the process (*Ptk2, Cacna2d1, Sh3glb1*). Taken together, these observations argue that microexon regulation by *Srrm3* modulates several processes and pathways that affect the regulation of insulin secretion in beta cells.

### *Srrm3* is necessary for microexon inclusion and proper insulin secretion regulation in mouse islets

To investigate the physiological role of *Srrm3*-regulated IsletMICs in islet function and glucose metabolism *in vivo*, we took advantage of a recently published constitutive *Srrm3* gene-trapped mouse line^18^. We bred mice with homozygous or heterozygous deletions, and analyzed islets at 10-12 weeks to assess alternative splicing by RT-PCR and RNA-seq. Consistent with the observations made in beta cell line knockdowns, the majority of IsletMICs were down-regulated in *Srrm3* homozygous mutant (Srrm3 −/−) islets (Fig. 4A and Extended Data Fig. 4A,B), and the lack of *Srrm3* had a preferential impact on IsletMICs compared to other alternative exons (Fig. 4B). A milder, but highly significant down-regulation of IsletMICs was also observed in heterozygous mutant mice (Srrm3 +/−) (Fig. 4A and Extended Data Fig. 4B-D). To investigate the effect of the loss of *Srrm3* on islet function, we performed static secretion experiments following stimulation with high concentration of glucose or an amino acid mix. In line with the results in cell lines (Fig. 3A,B), Srrm3 −/− islets displayed increased stimulated insulin secretion following exposure to high glucose (Fig. 4C) and to amino acids (Fig. 4D). Total cellular insulin was decreased in Srrm3 −/− islets (Extended Data Fig. 4E), while insulin mRNA levels were unaltered (Extended Data Fig. 4F), suggesting a depletion of intracellular insulin stores. We also assessed glucagon secretion from alpha cells after exposing islets from high to low glucose concentrations. Resembling the effects on insulin secretion, Srrm3 −/− islets showed significantly increased glucagon release in response to 1 mM glucose (Fig. 4E), while the levels of glucagon expression were not affected (Extended Data Fig. 4G,H). In the case of Srrm3 +/−islets, a milder increase in insulin secretion was observed following exposure to high glucose (Fig. 4C) and to amino acids (Fig. 4D), but, contrary to Srrm3 −/−, no differences in glucagon secretion were detected (Fig. 4E). Altogether, these findings indicate that loss of *Srrm3* leads to defective islet hormone secretion.

**Figure 4.**
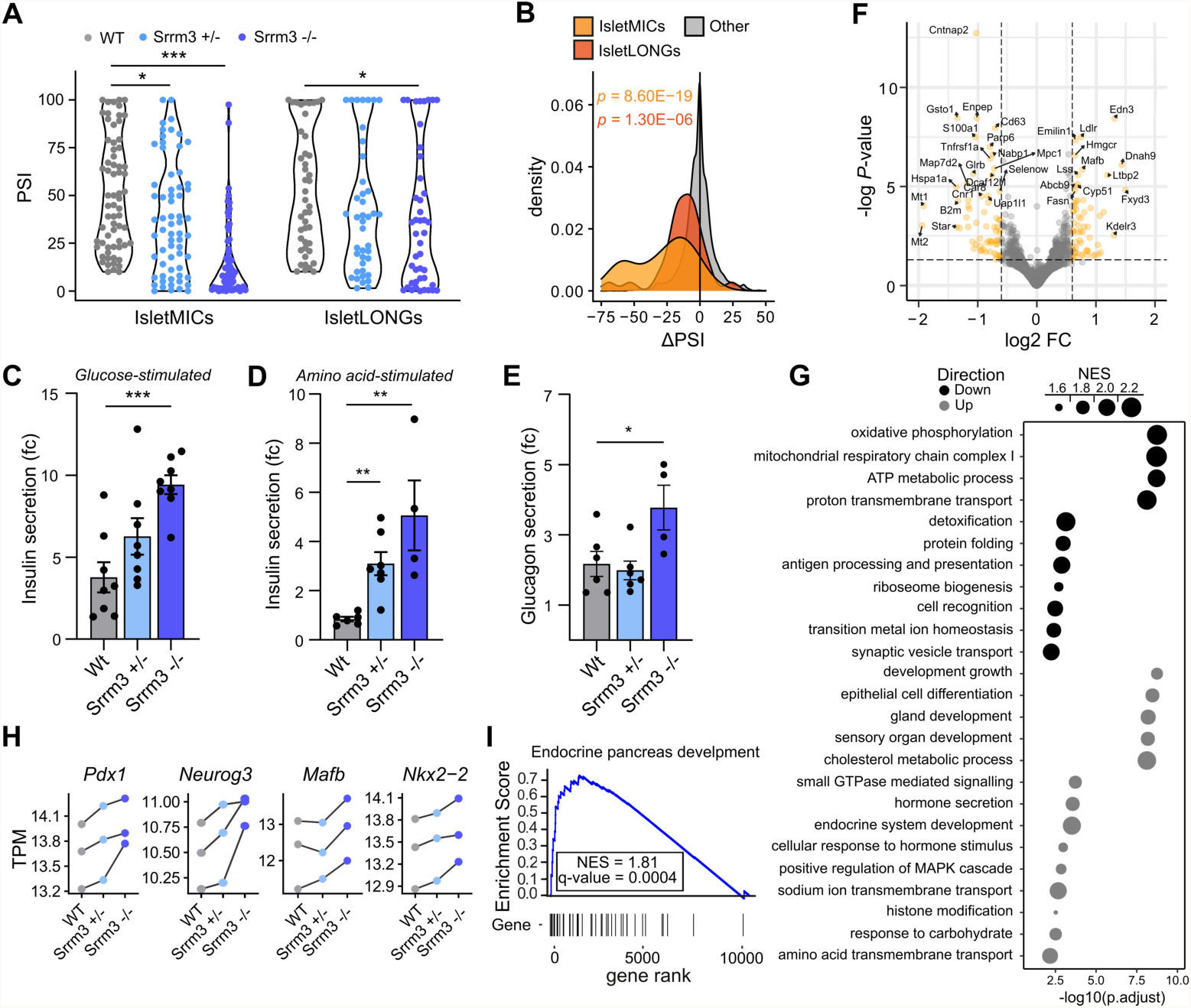
*Srrm3* depletion in mouse islets causes IsletMIC downregulation and increased stimulated insulin release. **A)** Exon inclusion levels in islets from Srrm3 mutant vs wild type mice for IsletMIC and IsletLONG programs. PSI values are the mean of three mice islet preparations. **B)** Density plots for ΔPSI distributions in Srrm3 −/− islets of IsletMICs, IsletLONGs and other alternative exons. P-values were obtained from two-sided Wilcoxon test comparing the distributions of IsletMICs (orange) or IsletLONGs (red) against other alternative exons. **C)** Fold change in glucose-stimulated insulin secretion following sequential stimulation from 2.8 mM to 20 mM glucose in cultured islets of each genotype (*n* = 8, mean ± s.e.m). **D)** Fold change insulin secretion following sequential stimulation from 0 mM to 12 mM of a mix of six amino acids (*n* = 4-6, mean ± s.e.m). **E)** Fold change in glucagon secretion following sequential stimulation from 20 mM to 1 mM glucose (*n* = 4-6, mean ± s.e.m). **F)** Volcano plot showing differentially expressed genes in Srrm3 −/− islets (|log2FC| ≥ 0.25) (*n* = 3). **G)** Representative enriched Biological Process GO terms for genes downregulated (black) or upregulated (grey) in islets from Srrm3 −/− mutant mice. **H)** mRNA expression levels for several transcription factors involved in endocrine pancreas development. Lines connect data obtained for different mice from the same litter/batch. **I)** Gene set enrichment analysis of endocrine pancreas development genes showing significant up-regulation of the pathway in Srrm3 −/− islets. Statistical comparisons in (A) and (C-E): *, p < 0.05; **, p < 0.01 and ***, p < 0.001 versus WT; unpaired t test.

To shed light into the molecular basis of these phenotypes, we next performed a differential gene expression analysis of *Srrm3* mutant and WT islets. A total of 193 and 286 genes were down-and up-regulated, respectively, in islets from Srrm3 −/− mice relative to WT (padj < 0.05, |log2FC| ≥ 0.25) (Fig. 4F). The regulated genes showed no significant overlap with genes harboring IsletMICs or IsletLONGs (Extended Data Fig. 4I,J), indicating that changes in gene expression are not directly caused by misregulation of islet-enriched exons. Down-regulated genes were mainly enriched in mitochondrial genes related to oxidative phosphorylation and ATP production by cellular respiration (Fig. 4G, Supplementary Table 4), in apparent contradiction with the increased insulin release observed both *ex vivo* and *in vivo*. Nevertheless, several pathways that play a positive role in insulin secretion were enriched among the up-regulated genes, including small GTPase-mediated signaling, hormone secretion, response to carbohydrates and amino acid transmembrane transport (Fig. 4G, Supplementary Table 4). These data suggest that altered nutrient sensing and an augmented secretory response compensate the decrease in mitochondrial respiration, resulting in net increase of insulin secretion. Remarkably, up-regulated genes were also enriched in GO terms related to organ morphogenesis and differentiation of epithelial cells, including endocrine system development (Fig. 4G, Supplementary Table 4). These up-regulated genes include several transcription factors with key roles in islet cell differentiation, such as *Pdx1, Neurog3, Mafb* and *Nkx2*.*2* (Fig. 4H). Moreover, Gene Set Enrichment Analysis (GSEA) revealed a general up-regulation of the endocrine pancreas development pathway in Srrm3 −/− islets (Fig. 4I), suggesting that loss of *Srrm3* can affect islet morphogenesis.

### Loss of *Srrm3* disrupts islet cell composition and architecture

To further investigate how *Srrm3* loss affected islet development and cell identity, we performed histological analysis of mice pancreata at 4 weeks after birth. *Srrm3* mutant islets presented alterations in cell type composition and architecture (Fig. 5A). In particular, Srrm3 −/− islets showed a higher proportion of alpha cells than WT islets (Fig. 5B), with decreased stereotypical location at the islet periphery (Fig. 5C). Moreover, Srrm3 −/− islets presented a significant increase in the number of polyhormonal cells co-expressing both insulin and glucagon (Fig. 5D). Islets from Srrm3 +/−mice showed similar but milder alterations (Fig. 5A-D). Next, to determine whether these alterations occurred during embryonic development or were acquired postnatally, we analyzed pancreata from newborn mice. Similar to 4-week-old mice, neonatal islets of mutant mice showed signs of altered endocrine cell differentiation, with an increased number of polyhormonal cells in Srrm3 −/− (Extended Data Fig. 5C). Additionally, Srrm3 −/− pancreata showed alterations in the clustering of endocrine cells, presenting a higher proportion of disorganized islet structures and dispersed endocrine cells (Extended Data Fig. 5D). Taken together, these findings indicate that loss of *Srrm3* alters normal islet morphogenesis during pancreas development.

**Figure 5.**
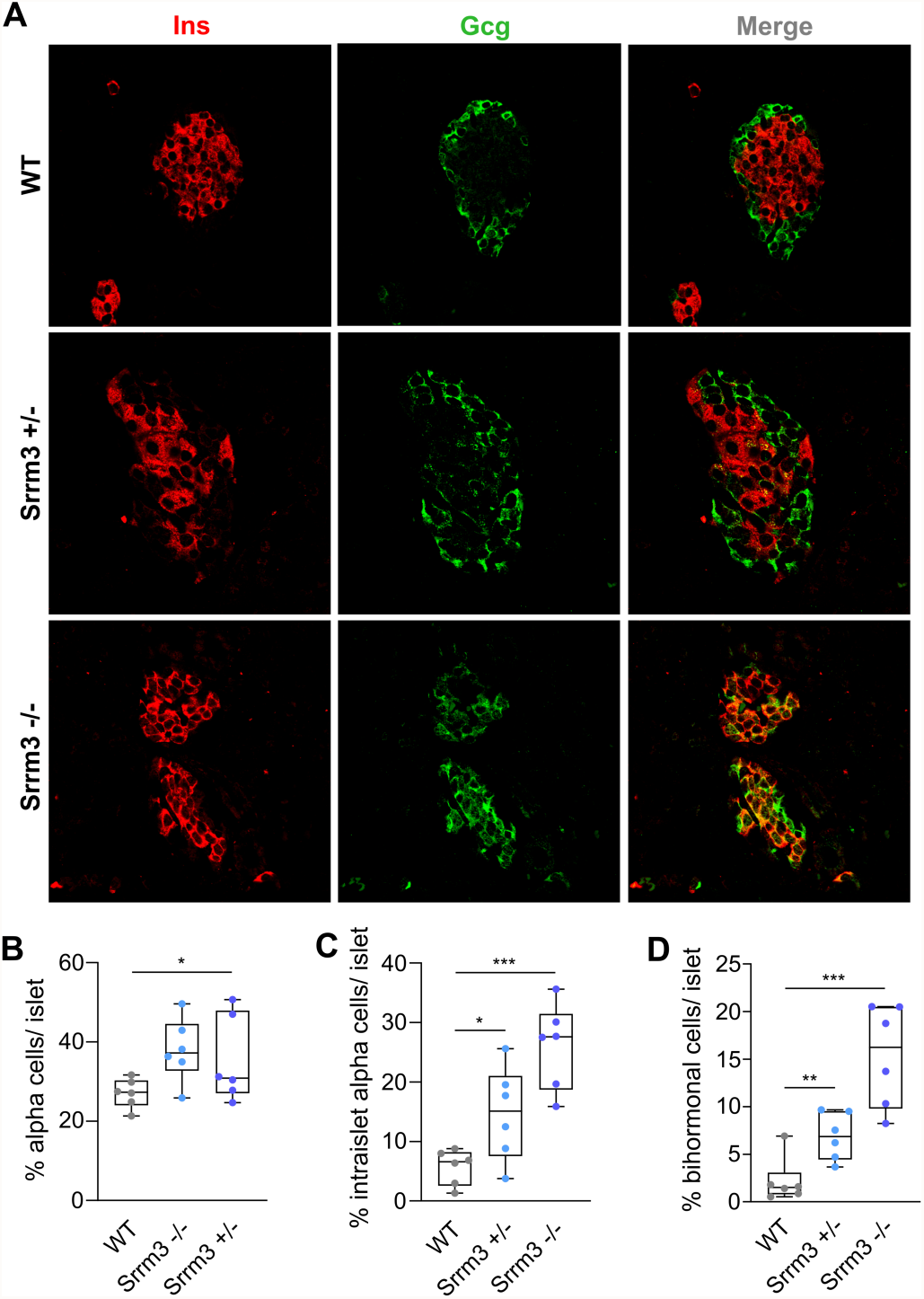
*Srrm3* depletion disrupts islet cell composition and architecture. **A)** Representative immunofluorescence images of islets from WT and *Srrm3* mutant mice stained for insulin (Ins; red) and glucagon (Gcg; green) at 4 weeks of age. **B)** Percent of alpha cells in WT and *Srrm3* mutant islets. **C)** Percent of alpha cells outside the islet periphery, indicating disrupted islet architecture in *Srrm3* mutant mice. **D**) Percent of insulin and glucagon double positive cells in WT and *Srrm3* mutant mice. All data are presented as mean ± s.e.m (*n* = average of 9-12 islets from 6 mice per genotype including males and females). Statistical comparisons: *, p < 0.05; **, p < 0.01 and ***, p < 0.001 versus WT; unpaired t test.

### *Srrm3* depletion leads to hyperinsulinemic hypoglycemia in mice

We next investigated if the functional and histological alterations observed in the islets of *Srrm3* mutant mice impacted glycemic control *in vivo*. We first performed blood glucose measurements in adult mice fed *ad libitum* and found that Srrm3 −/− mice presented drastically decreased glycemia compared to WT and Srrm3 +/−littermates (Fig 6A). Next, we measured blood glucose after 4h of fasting and 30 min after food intake (postprandial). Srrm3 −/− mice presented strong hypoglycemia in both fasting and postprandial conditions, while glycemia in Srrm3 +/−mice was normal at fasting but significantly reduced in postprandial conditions (Fig. 6B). In order to test whether hypoglycemia in Srrm3 −/− mice was associated with excessive insulin secretion from pancreatic islets, we measured insulin plasma levels at fasting and postprandial conditions. While insulin was undetectable at fasting in most samples for all genotypes, Srrm3 −/− mice showed a significant increase in plasma insulin and insulin/glucose ratio at 30 min postprandial compared to WT mice (Fig. 6C,D). We also found that Srrm3 −/− mice showed reduced glucagon levels at fasting (Fig. 6E), potentially due to insulin-induced inhibition of glucagon secretion. No major differences between male and female mice were observed in either glycemia or hormone plasma levels (Extended Data Fig. 6A–F). Taken together, these findings are compatible with hyperinsulinemic hypoglycemia induced by depletion of *Srrm3* in homozygous mutant mice.

**Figure 6.**
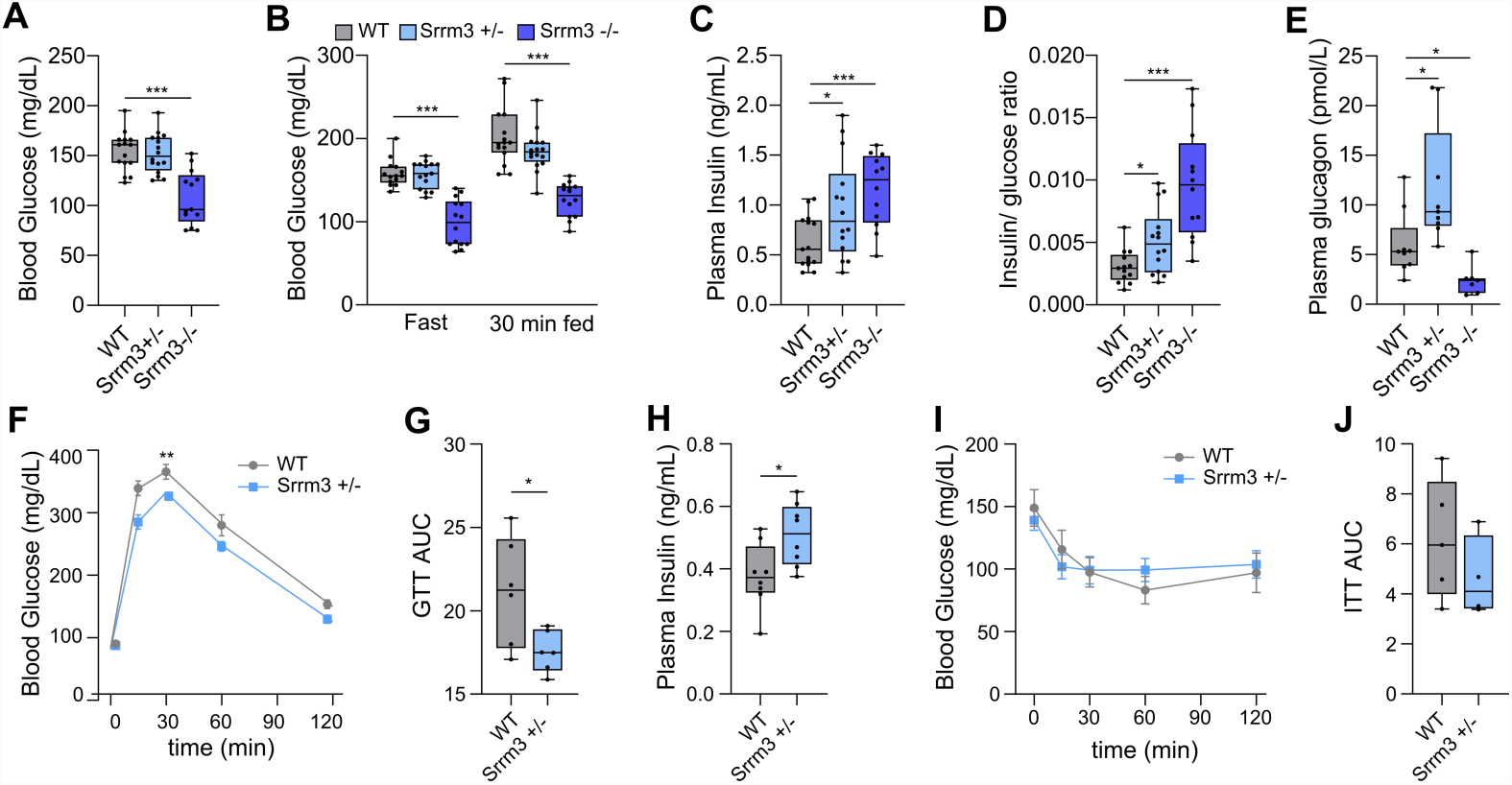
*Srrm3* mutant mice display hypoglycemic hyperinsulinemia. **A)** Random blood glucose measurements in *Srrm3* mutant and wild type mice fed ad libitum (*n* = 13-15). **B)** Blood glucose levels following a fast and fed test (*n* = 14-15). Glucose was measured at the end of a 4h fast and at 30 min postprandial (30 min fed). **C**,**D)** Insulin plasma levels (C) and ratio between plasma insulin and blood glucose (D) at 30 min postprandial (*n* = 12-15). **E)** Plasma glucagon at 4h fasting (*n* = 7-9). **F)** Intraperitoneal glucose tolerance test in WT and Srrm3 +/−mice, showing blood glucose levels at different time points (*n* = 6, mean ± s.e.m). **G)** Area under the curve (AUC) for each mouse shown in (F). **H)** Plasma insulin at 15 min after intraperitoneal glucose injection. **I)** Insulin tolerance test in WT and Srrm3 +/− mice, showing blood glucose levels at different time points (*n* = 5, mean ± s.e.m). **J)** Area under the curve (AUC) for each mouse shown in (I). Statistical comparisons: *, p < 0.05; **, p < 0.01 and ***, p < 0.001 versus WT; unpaired t test. Measurements shown in A-E were performed using both male and female mice, while tolerance test shown in F-J were performed only in males.

Because Srrm3 −/− mice displayed reduced body weight and neurological defects (Nakano et al. 2019), which may also contribute to altered glycemia, we next focused on Srrm3 +/− mice, which showed no differences in growth or evident neurological defects but exhibited functional and histological alterations in islets. Intraperitoneal glucose tolerance test (IPGTT) showed mild but significantly increased glucose tolerance in Srrm3 +/− compared to WT mice (Fig. 6F,G). This increase was associated with augmented plasma insulin levels at 15 min after glucose injection (Fig. 6H). On the other hand, intraperitoneal insulin tolerance test (ITT) showed no significant differences in sensitivity to exogenously administered insulin between Srrm3 +/− and WT mice (Fig. 6I,J). Collectively, these observations are compatible with the concept that loss of *Srrm3* impairs glucose homeostasis due to inappropriate insulin secretion.

### *SRRM3* harbors genetic variants associated with fasting glucose and T2D risk

We next explored possible associations between human genetic variants in the *SRRM3* locus and either T2D susceptibility or glycemic traits. We found a suggestive association between T2D risk and variants downstream of the *SRRM3* gene in the Finnish population (Finngen study, r5.finngen.fi) (rs7459185; β = −0.0565, *P* = 2.1 × 10^−6^) (Fig. 7A), although this association was not observed in other human populations (^25^ and CMDKP database, hugeamp.org). On the other hand, the analysis of a trans-ancestral large-scale meta-analysis of glycemic traits^26^ revealed that rs67070387, a polymorphism within *SRRM3* intron 1, is strongly associated with increased fasting glucose (FG) levels (β = 0.0233, *P* = 7.94 x 10^−12^) (Extended Data Fig. 6A). Additional variants in *SRRM3* and in the nearby gene *STYXL1* associated with the same trait are also reported in the CMDKP database (hugeamp.org) (Extended Data Fig. 7A), although credible set analysis did not confidently identify the likely causal variant^26^. Next, we investigated whether these genetic variants overlap with previously annotated regulatory elements in human islets. The variant with suggestive T2D association, rs7459185, and its associated linkage disequilibrium block are located in a linear enhancer cluster active in human islets that directly interacts with *SRRM3* promoter according to capture Hi-C and Hi-ChIP chromatin analyses (Fig. 7B^4^). On the other hand, the FG variant rs67070387 maps to an intronic region near the *SRRM3* promoter (Fig. 7B) that is annotated as a weak enhancer in the ENCODE database. This variant is predicted to disrupt a binding site for *FOXA2* (Extended Data Fig. 6B), a transcription factor that participates in the regulation of the insulin secretory pathway^27^.

**Figure 7.**
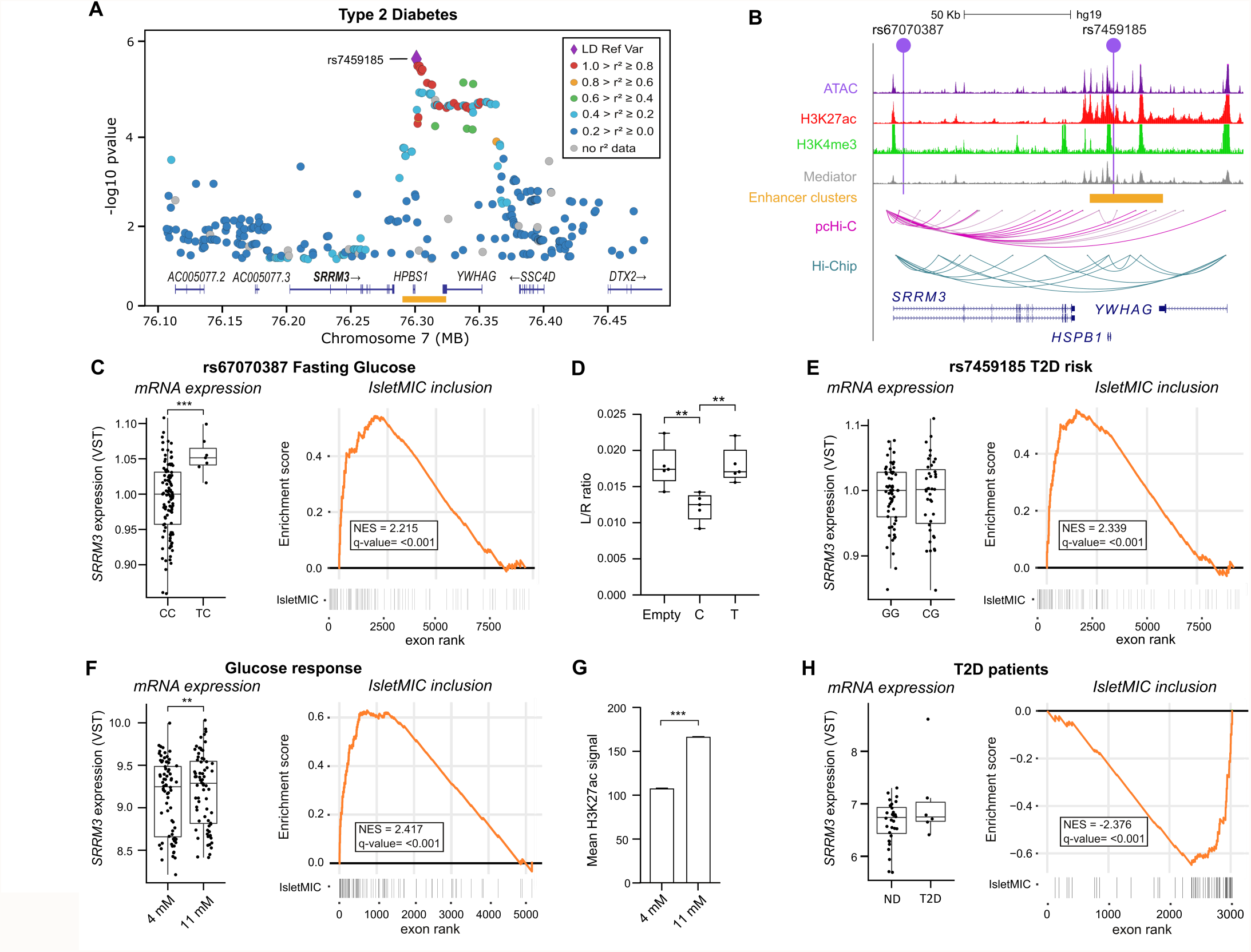
The *SRRM3* locus responds to glucose and harbors genetic variants associated with fasting glucose and type 2 diabetes risk. **A)** Regional plot of T2D GWAS variants from the FinnGen study (r5.finngen.fi/gene/SRRM3). An islet-specific enhancer cluster regulating *SRRM3* is shown in orange. **B)** Integrative map of the *SRRM3* locus in human islets showing chromatin state profiles and pcHi-C interactions between clustered enhancers and the gene promoter ^4^. **C)** Normalized *SRRM3* expression and microexons inclusion (measured by alternative splicing set enrichment analysis) in human islets from two independent datasets according to FG rs67070387 genotype (*n* = 7-103). **D)** Luciferase assay in EndoC-βH1 cells following transfection of a control reporter vector or carrying the genomic sequence surrounding rs67070387 with one or the other allele (*n* = 5). **, p < 0.01 versus indicated by unpaired t test. **E)** Normalized *SRRM3* expression and IsletMIC inclusion in human islets from two independent datasets according to T2D rs7459185 genotype (*n* = 39-60). **F)** *SRRM3* expression and IsletMIC inclusion following exposure of human islets to high/low glucose (*n* = 72). **G)** Mean H3K27ac signals of the *SRRM3* enhancer cluster in human islets following exposure to low and high glucose. **H)** *SRRM3* expression and IsletMIC inclusion in human islets from non-diabetic (ND) and type 2 diabetes (T2D) individuals (*n* = 6-34) from^28^.

To test whether these variants are associated with transcriptional variation in *SRRM3* expression and/or IsletMICs inclusion, we analyzed human islet RNA-seq data from 110 genotyped non-diabetic donors from two different sources^28, 29^. Despite the low allele frequency (Extended Data Fig. 7B), we found a significant association between the FG variant rs67070387 and increased *SRRM3* expression in *cis*, as well as higher microexon inclusion in *trans* (Fig. 7C). To validate this association, we cloned the 500-nt region containing the rs67070387 SNP in a vector containing a luciferase cDNA reporter under a minimal promoter and performed luciferase assays in EndoC-βH1 and INS-1E cells. While the most common variant reduced basal luciferase expression, suggesting that it harbors a transcriptional repressive activity, the rare FG variant associated with increased fasting glucose abolished this effect, displaying higher luciferase expression in both human (Fig. 7D) and rat (Extended Data Fig. 7C) cells. This is in line with the differences in *SRRM3* expression between the two genotypes observed in the human islet data (Fig. 7C). In the case of rs7459185, although no differences in *SRRM3* expression were found between genotypes under these conditions (Fig. 7E, left panel), the genotype associated with decreased T2D risk showed a significantly increased inclusion of IsletMICs (Fig. 7E, right panel).

We next investigated whether genetic variants in IsletMICs or their flanking intronic regions were associated with fasting glucose or increased risk of T2D. To that end, we leveraged quantile-quantile (QQ) plots to compare the distribution of trait association p-values against the expected null distribution for variants located between 500 nt upstream and downstream of IsletMICs. While no major inflation was found for T2D (Extended Data Fig. 7D, left panel), we observed a suggestive trend of lower FG association p-values at microexon variants (Extended Data Fig. 7D, right panel). Although our results are limited due to the relatively low number of IsletMICs under study, this finding raises the possibility that variants affecting splicing regulatory sequences and microexon inclusion levels may impact islet function and glycemic control.

### IsletMICs respond to glucose and are misregulated in islets of T2D patients

To further substantiate the physiological relevance of the IsletMIC program, we next explored whether IsletMICs and/or *SRRM3* could be dynamically regulated in response to extracellular signals. First, we analyzed RNA-seq data of human islets exposed to moderately low (4 mM) or high (11 mM) glucose for 72 h^4^, and found that both *SRRM3* expression and microexon inclusion were significantly increased under high glucose (Fig. 7F). Interestingly, these changes correlated with an increase of the open chromatin H3K27ac signal on the *SRRM3* enhancer cluster (Fig. 7G), arguing that the IsletMIC program is directly induced by glucose.

Finally, we assessed whether the IsletMIC program is misregulated in islets from individuals with T2D. RNA-seq analysis from two independent studies^28,30^ (revealed that, while *SRRM3* mRNA levels were not significantly altered (Fig. 7H and Extended Data Fig. 7E left panels), IsletMIC inclusion was strongly down-regulated in T2D islets (Fig. 7H and Extended Data Fig. 7E right panels). A similar pattern was also found in islets from diabetes-susceptible NZO mice following a diabetogenic stimulus (Extended Data Fig. 7F)^31^. Importantly, down-regulation of IsletMICs was unlikely to be due to beta cell loss, as we also observed it in individuals with impaired glucose tolerance (Extended Data Fig. 7G,H) and in single beta cells form T2D individuals (Extended Data Fig. 7I). Collectively, our findings are consistent with the concept that the IsletMIC program is dynamically regulated under physiological and pathological conditions and argue that human genetic variation affecting islet-enriched regulation of *SRRM3* and overall IsletMIC inclusion levels has an impact on human glucose homeostasis.

## Discussion

While alternative splicing is known to play crucial roles in the development and function of a variety of cell and tissue types^9^, its impact on endocrine pancreas and its physiological relevance in the context of glycemic control remains largely unknown. Here, we identify a novel conserved program of islet-enriched alternative microexons and provide diverse *in vitro, in vivo* and genetic evidence that it plays important roles in maintaining islet function, globally impacting the regulation of glucose homeostasis and T2D susceptibility.

We show that IsletMICs represent a subset of the broader microexon program previously described as neural-enriched and regulated by the neural-specific expression of the splicing factor *SRRM4*^13^. Our work reveals that this previous paradigm is in fact part of a more complex regulatory design, whereby two nested tissue-enriched programs are generated by the tiered expression of a single regulatory activity (provided by SRRM3 and/or SRRM4) and the different intrinsic sensitivities of its target microexons. By expressing relatively low levels of *SRRM3*, in contrast to the higher levels of both *SRRM3* and *SRRM4* in neurons, islet cells recruited only the most sensitive microexons among the larger neural microexon program. Such nested regulation likely reflects the deployment of common biological processes for the function of both pancreatic endocrine cells and neurons. In line with this hypothesis, IsletMICs are strongly enriched for processes involved in secretory exocytosis, including vesicle transport, ion channel activity, vesicle priming or cytoskeleton organization, processes that are shared and essential for hormone and neurotransmitter release. These findings are in line with previous studies uncovering the expression of other neural-enriched RNA binding proteins, such as Nova1 and Rbfox1/2, in beta cells^10, 11^. Moreover, a shared regulation of alternative splicing fits with the general observation that mRNA expression and chromatin marks of islet cells are similar to those of neurons despite their different embryological origin^32^ These patterns are thus consistent with the hypothesis that pancreatic endocrine cells have co-opted regulatory signatures from neurons^33, 34^.

In line with a functional role of IsletMICs in vesicle exocytosis, we show that *SRRM3* depletion in human and rat beta cell lines leads to increased glucose-stimulated insulin secretion. Our data suggest that *SRRM3* impacts several processes involved in insulin secretion in beta cells, including energy metabolism and cytoskeletal reorganization. In agreement with this, individual down-regulation of IsletMICs included in genes involved in different biological processes such as vesicle formation (*SH3GLB1*), cytoskeleton reorganization (*PTK2*), calcium signaling (*CACNA2D1*) and granule exocytosis (*CASK*) was sufficient to alter glucose-stimulated secretion. Similar to cell lines, islets from *Srrm3* mutant mice showed an inappropriate increase in insulin secretion. At the physiological organismal level, this increase correlated with higher postprandial plasma insulin levels and fasting hypoglycemia in homozygous mice, suggesting that the lack of microexon inclusion in islets leads to hyperinsulinemic hypoglycemia. Importantly, heterozygous mice, which show no growth or neurological defects compared to their homozygous littermates^18^, presented a similar but milder phenotype characterized by increased glucose tolerance. Although future work using tissue-specific ablation of *Srrm3* will be required to determine the precise impact of islet microexons *in vivo*, our data collectively support an important role for the *Srrm3*-regulated islet splicing program in insulin secretion, impacting various of the underlying processes and possibly also islet cell differentiation and morphogenesis.

Several lines of evidence suggest that *SRRM3* has also relevance for human glucose homeostasis and diabetes pathophysiology. Firstly, we found genetic variants in both *SRRM3* and microexons associated with fasting glucose or T2D risk. Secondly, exposure of human islets to mildly elevated glucose concentrations led to increased *SRRM3* expression and IsletMIC inclusion, suggesting an adaptive mechanism to modulate insulin secretion in a setting that has the potential to enhance secretion. Finally, we also observed that IsletMICs are broadly down-regulated in islets from T2D patients, an effect also recently reported in a diabetic mouse model^31^. Altogether, these data suggest that changes in the activity of SRRM3 and its microexon targets affect islet function and glucose homeostasis, potentially contributing to T2D predisposition. There is currently a broad range of strategies to manipulate RNA splicing therapeutically ^35^. Therefore, future research on the molecular function of IsletMICs and other islet-enriched exons and their role in islet dysfunction might provide valuable knowledge not only for islet biology but also for the development of novel therapies for T2D based on splicing modulation.

## Acknowledgements

We thank Botond Banfi for kindly sharing the Srrm3 gene-trapped mouse line with us; Miguel Ángel Maestro for excellent technical advice on multiple protocols related to the study of *Srrm3* mutant mice; Jon Permanyer and Cristina Rodriguez for help with mouse genotyping; Diego Balboa, Irene Miguel-Escalada and Edgar Bernardo, as well as members of the Irimia and Valcárcel groups for constant scientific discussion; André Gohr for assistance on bioinformatic analyses; Stephen Taylor for kindly sharing the HeLa Flp-In T-Rex cell line with us; and CRG Genomics and Advanced Light Microscopy Units for the RNA sequencing and microscopy services.

## METHODS

### Definition of islet-enriched alternative exons

Inclusion tables for human and mouse were downloaded from VastDB website (vastdb.crg.eu)^8^, together with additional islet and beta and alpha cell RNA-seq samples downloaded from the NCBI Short Read Archive (SRA) (Supplementary Table 1). We also downloaded available RNA-seq data from SRA for multiple differentiated tissues and embryonic samples for *Rattus norvegicus* and created a similar inclusion table for all alternative splicing events using *vast-tools* v2.5.1 (see below), which is also accessible in VastDB (Supplementary Table 1).

To define islet-enriched and neural-enriched exon programs we first parsed these RNA-seq datasets to identify tissue-specific exons using the script *Get_Tissue_Specific_AS*.*pl* script^36^ (https://github.com/vastdb-pastdb/pastdb), requiring the following cut-offs: i) absolute difference in the average exon inclusion level (using the percent-spliced-in metric, PSI) between the target tissue and the averages across other tissues of |ΔPSI| > 15 (--min_dPSI 15), ii) global |ΔPSI| > 25, i.e., difference between target tissue inclusion average and the average of all other tissues combined (-- min_dPSI_glob 25), iii) a valid average PSI value in at least N = 5 tissues (--N_groups 5), and iv) sufficient read coverage in at least n = 2 samples per valid tissue group (--min_rep 2), i.e., score VLOW or higher as provided by *vast-tools*^8^. For each species, a config file with the tissue groups for each RNA-seq sample was provided as input (Supplementary Table 5). To better capture all microexons with biased inclusion in certain tissue groups, we excluded from the tissues with known or expected partial overlap of microexon inclusion from comparisons in those tissues (for example, neural, endocrine pancreas, muscle and heart), as provided in the column EXCLUDED in the config files (Supplementary Table 5). Next, for those exons defined as up-regulated in neural and/or endocrine pancreas by this analysis, we added two additional steps. First, if the difference between the average PSI in neural/endocrine pancreas samples and in other tissues (ΔPSI, as provided by *Get_Tissue_Specific_AS*.*pl* using the --test_tis option) was higher than 15, the exon was further considered as neural/islet enriched. Second, enriched exons whose average PSI in other tissues was higher than 25 were discarded.

### Gene Ontology and GWAS gene enrichment analyses

Gene Ontology (GO) analyses of IsletMICs were performed gprofiler2 R package (Kolberg et al., 2020), with custom annotations and gene backgrounds. Human GO annotations were obtained from^37^. For GO analysis of human IsletMICs, we provided as a custom gene background all genes harboring at least one splicing event of any type from *vast-tools* passing the same filtering criteria required in the analysis described above using the *Get_Tissue_Specific_AS*.*pl* script for the islet-enriched exons (the list of such events was obtained using the option --test_tis). We identified 106 GO terms for Biological Process passing FDR ≤ 5%. We then manually curated GO terms and collapsed them into 10 generalized functions important for the endocrine pancreas, taking the FDR value of the largest terms within the new group.

To test the overlap between genes containing islet-enriched exons and GWAS variants of interest, we first downloaded from the NHGRI-EBI GWAS Catalog^38^ all unique genes harboring significant GWAS variants associated with disease traits matching “Type 1 diabetes”, “Type 2 diabetes” or “Glycemic traits” (download date 2021-12-05; https://www.ebi.ac.uk/gwas/) (Supplementary Table 6). To test the overlap with genes containing islet-enriched exons, we first discarded from these sets the genes that did not contain at least one splicing event that passed the filtering criteria used to define islet-enriched exons (see above). The significance of the overlap between each of these sets and genes harboring IsletMICs/IsletLONGs was assessed using two-sided Fisher’s Exact tests, with the same gene background as per GO analyses.

### Single Cell RNA-Seq analysis

Publicly available islet single cell RNA-seq samples from multiple donors, studies and technologies were used to identify cell specific expression and splicing profiles. Reads from plate-based methods (SRP075377^39^, SRP076307^40^, ERP017126^41^ and SRP075970^42^), which provide full-transcript information, were mapped to human transcriptome (Ensembl release 104) using Salmon^43^ (version 1.4.0) for gene quantification, generating a gene expression table per study. In addition to these samples, we used the gene counts from Drop-Seq technology for an additional study (SRP111557)^44^. All gene counts were integrated (top 3000 variable features) after normalization (top 5000 variable features) with Variance Stabilizing Transformation (VST) from sctransform package ^45^ into a single table, and then the Uniform Manifold Approximation and Projection technique (on the first 30 PCs) was used for dimension reduction using Seurat R package^46^. Cell clustering was calculated using a resolution of 0.3. We annotated each cluster based on the cell type with the highest proportion per cluster using the CellID pipeline (50 dimensions using 200 features)^47^. Alternative splicing quantifications were obtained using *vast-tools* v2.5.1^8^ for hg38 with a pseudo-bulk approach, where FASTQ files were merged based on cell type. Inclusion levels, using the PSI metric, for all IsletMICs and IsletLONGs were pulled out of each cell type from non-T2D donors ignoring those with N or VLOW *vast-tools* read coverage score for comparing PSI among cell types.

### Evolutionary conservation and open reading frame impact prediction

To investigate the evolutionary conservation of human IsletMICs and IsletLONGs in rodents (Fig. 1D), we performed the following steps. First, genome-conservation was assessed as previously described^13^. Briefly, exon coordinates from human exons (hg38) were lifted to the mouse (mm10) or rat (rn6) genomes with the *liftOver* tool. Then, lifted exon coordinates in the target rodent species were matched with the exon coordinates from the *vast-tools* mm10 or rn6 annotation. Then, to investigate islet-enriched regulatory conservation, we first obtained ΔPSI values (average PSI in islets - average PSI in other tissues) from the *Get_Tissue_Specific_AS*.*pl* as described above (using the --test_tis option), but requiring only one replicate with sufficient read coverage per tissue group, to increase the number of tested exons (--min_rep 1). If the exon had sufficient read coverage, it was considered regulatory conserved if ΔPSI ≥ 15. Open reading frame impact predictions (Extended Data Fig. 1E) were done as previously described^13^, using the predictions provided in VastDB (v4; https://vastdb.crg.eu/wiki/Downloads).

### Alternative splicing and differential gene expression analyses

Alternative splicing was quantified using *vast-tools* v2.5.1^8^ for all RNA-seq datasets. FASTQ reads were aligned with *vast-tools align* employing the following VASTDB libraries: (i) human (hg38, vastdb.hs2.23.06.20), (ii) rat (rn6, vastdb.rno.23.06.20), and (iii) mouse (mm10, vastdb.mm2.23.06.20). The outputs of *vast-tools align* for each RNA-seq sample were then combined into a single table with all splicing events using *vast-tools combine* with default parameters. Differentially spliced exons upon *Srrm3* depletion were identified using *vast-tools compare*, with default parameters: |ΔPSI| > 15 between the means of the two compared groups, e.g. control vs SRRM3 siRNA (--min_dPSI 15), and a non-overlapping PSI distribution between two sample groups of at least 5 (--min_range 5).

Differential gene expression analysis was additionally performed for mouse islets (Figs. 4 and Extended Data Fig. 4). RNA-seq reads were mapped with Salmon^43^ for gene quantification to human or mouse transcriptome (Ensembl release 104). Raw gene counts were normalized with VST from DESeq2^48^ bioconductor package. Differential gene expression was assessed using DESeq2, after independent filtering, with a design which takes into account Genotype + Batch, and genes were considered differentially expressed if adjusted p-value was lower than 0.05 and absolute shrinked log fold change, calculated with apeglm method^49^, greater than 0.6. For the functional enrichment analysis (Fig. 4G), all genes assessed for differential expression (9,729) genes were sorted based on log2 fold change expression between *Srrm3* mutant and WT mice and used as input for a Gene Set Enrichment Analysis of GO with bioconductor package clusterProfiler^50^. Enriched terms shown in Fig. 4G were selected according to GO-terms Semantic Similarity Measures from GOSemSim^51^ based by p-value with a similarity cutoff of 0.5 and the “Jiang” method.

### Evaluation of microexon sensitivity to SRRM3

Stable cell lines overexpressing mouse *Srrm3* at three different levels were generated using HeLa Flp-In T-Rex cells (kindly provided by S. Taylor, University of Manchester) and Flp-In (Thermo Fisher Scientific) expression plasmids. Flp-In plasmids containing either GFP^19^ or 3x N-terminal-Flag tagged mouse *Srrm3* were generated using standard restriction and ligation (sequences of primers used for cloning are provided in Supplementary Table 7). HeLa Flp-In T-REx cells were transfected with 0,5 μg pcDNA5 vector expressing GFP^19^ or mouse *Srrm3* and 4,5 μg pOG44 vector in a 10 cm plate (10^6^ cells) using Lipofectamine 2000 (Invitrogen). Cells containing stably integrated constructs were selected using 150 μg/ml Hygromycin B until visible colonies were formed. Individual colonies were isolated and expanded. For the induction experiments (Fig. 2F), cells were plated at 200,000 cells per well of a 6-wells plate. The GFP expressing cells were grown without Doxycycline (Control) and Flag-Srrm3 expressing cells were treated with either no Doxycycline (low) or 0.6 (mid) and 16.7 (high) ng/ml Doxycycline for 24 hours. RNA was extracted using the RNeasy Plus Mini kit (Qiagen). RNA quality was checked using Bioanalyzer (Agilent), and strand-specific Illumina libraries were prepared and sequenced at the CRG Genomics Unit. Samples were sequenced on HiSeq2500 and 125 nts paired-end reads were generated. Number of reads and percentage of mapping to genome and transcriptome are described in Supplementary Table 1.

### Culture and Transfection of Rat INS-1E Cells and Human EndoC-βH1 Cells

Rat insulin-producing INS-1E cells, kindly provided by Dr. C. Wollheim (University of Geneva, Geneva, Switzerland), were cultured in RPMI 1640 GlutaMAX-I medium (Invitrogen) as described previously^52^. Human insulin-producing EndoC-βH1 cells, purchased from Univercell Biosolutions, were grown on plates coated with Matrigel/fibronectin (100 and 2 μg/ml, respectively, Sigma), and cultured in DMEM as described previously^53^. The small interfering RNAs (siRNA) targeting human and rat *Srrm3* and the antisense oligonucleotides (ASO) harboring 2′-O-methyl phosphorothioate modifications targeting individual microexons used in this study are described in Supplementary Table 6; Allstars Negative Control siRNA (Qiagen) was used as a negative control (siCTL). Transient transfection was performed using 30 nmol/L siRNA or 25 nmol/ L ASO and Lipofectamine RNAiMAX (Invitrogen).

### RNA analysis of beta cell lines and mouse islets

Total RNA was extracted using Maxwell RNAeasy kit (Promega) following the manufacturer’s instructions. For RNA-seq experiments, total RNA extraction was performed using RNeasy Mini Kit (Qiagen) following the manufacturer’s instructions. Reverse transcription (RT) was performed with 200 ng–1 μg of total RNA in the presence of 125 ng of random primers (Invitrogen), 50 pmol of oligo dT (Sigma) and 0.5 μL of Superscript III reverse transcriptase (Invitrogen) in a total volumen of 20 μL. Semiquantitative PCR reactions were carried out with 1–2 μL of cDNA and 0.2 μL of GoTaq DNA Polymerase (Promega) in a total reaction volume of 30 μL according to the manufacturer’s instructions. Typical parameters for the reactions were: 3 min at 94°C, 35 cycles of 30 sec at 94°C, 30 sec at 60°C, 30 sec at 72°C and a final incubation of 1 min at 72°C. Inclusion/exclusion patterns for selected alternative microexons were analyzed by RT-PCR followed by acrylamide electrophoresis and band densitometry using ImageJ. The primers were designed against flanking constitutive exons, allowing different splice variants to be distinguished based on fragment size. Quantitative PCR (qPCR) reactions were carried out with 4 μL of diluted cDNA, 0.4 μM of each primer, and 5 μL of Power SYBR Green PCR master mix (Applied Biosystems) in a total reaction volume of 10 μL. Technical replicates for each experimental sample were analyzed in a ViiA 7 Real-Time PCR system (Applied Biosystems) using the standard curve method. Relative expressions were calculated using the quantity mean normalized by actin or Gapdh. All primers used in this study are described in Supplementary Table 7. For in-house RNA sequencing, RNA quality for three independent experiments, each composed of a trio of sex-matched WT, heterozygous and homozygous mutant siblings, was assessed using Bioanalyzer (Agilent) and strand-specific Illumina libraries were prepared and sequenced at CRG Genomics Unit. Samples were sequenced on HiSeq2500 and 125 nts paired-end reads were generated. Number of reads and percentage of mapping to genome and transcriptome are described in Supplementary Table 1.

### Quantification of insulin secretion from INS-1E and EndoC-βH1 cells

To assess insulin secretion in INS-1E, cells were first pre-incubated for 1 h in glucose-free RPMI 1640 GlutaMAX-I medium (Life Technologies, Inc.), followed by incubation with Krebs-Ringer buffer (KRB) (140 mM NaCl, 3.6 mM KCl, 0.5 mM NaH2PO4, 2 mM NaHCO3, 1.5 mM CaCl2, 0.5 mM MgSO4, 10 mM HEPES, 0.25% BSA, pH = 7.4) for 30 min. Next, cells were exposed to 1.7 mM glucose (low) followed by sequential exposure to 17 mM glucose (high) with or without 20 μM forskolin or 100 μM tolbutamide during 30 min. Dynamic insulin secretion measurements was performed by collecting supernatants every 4 min after 17 mM glucose stimulation. To assess insulin secretion in EndoC-BH1, cells were first pre-incubated with culture medium containing 2.8 mmol/L glucose for 18 h. Next, cells were incubated in KRB for 1 h and sequentially stimulated with 1 mM glucose (low), 20 mM glucose (high), or 20 mM glucose plus 20 μM forskolin for 40 min. Supernatants were collected to assess insulin secretion and cells were sonicated in an alcohol-acid solution to measure total insulin content using Rat and Human Insulin ELISA Kits (Mercodia).

### Actin and tubulin cytoskeleton measurements

INS-1E cells were cultured on glass slides coated with Matrigel-fibronectin (100 and 2 μg/ml, respectively; Sigma) and transfected with control or *Srrm3* siRNAs as previously described. Following the same protocol used to assess glucose-stimulated insulin secretion, cells were pre-incubated for 1 h in glucose-free medium followed by 30-min incubation with KBR, and 30-min incubation with KBR 1.7 mM glucose. Cells were then exposed to 17 mM glucose for 5 or 20 minutes and immediately after fixed in 4% paraformaldehyde for 10 min. Cells were then washed and permeabilized with Triton X-100 0.25% for 3 min and blocked using 3% BSA. Next, cells were incubated at room temperature in sequential order with deoxyribonuclease I-Alexa Fluor 488 conjugate (Life Technologies, Inc.) and Atto 550-phalloidin (Sigma) to specifically stain G-actin (globular) and F-actin (filaments), respectively, or with an antibody against β-tubulin (1:500; Sigma). Nuclei were stained with Hoechst 33342. Slides were imaged by confocal microscopy using a Leica TCS SP5 and images processed by LAS X (Leica). ImageJ software was used to measure actin-rich filopodia and to quantify the intensity ratio between F-actin and G-actin, or β-tubulin.

### Measurements of intracellular ATP levels

To monitoring intracellular ATP levels under low and high glucose conditions, INS-1E cells were co-transfected with control or *Srrm3* siRNAs together with a vector encoding for a cytoplasmic ATP FRET probe^22^. Cells were transfected in 24 well plates, at a density of 300,000 cells per well, with 200 ng of vector and 30 nM of siRNAs. After 48 h, cells underwent the same protocol used to assess glucose-stimulated insulin secretion. Briefly, cells were pre-incubated in glucose-free medium for 1 h followed by KBR for 30 min. Next cells, were exposed to KRB 1.7 mM glucose for 30 min, followed by stimulation with KBR 17 mM glucose. Individual cells were imaged at 1.7 mM glucose (time 0) and at different time points following 17 mM glucose stimulation using a Leica TCS SP5 confocal microscopy and LAS X software (Leica,). FRET indexes per cell and time point were obtained by analyzing images with the ImageJ Plug-in “FRET and co-localization analyzer” (rsb.info.nih.gov/ij/plugins/fret-analyzer.html).

### *Srrm3* knock out mouse

C57BL/6 *Srrm3* gene-trapped mice were kindly provided by Botond Banfi (University of Iowa) ^18^. The gene trap is located in the third intron of *Srrm3*, and it encodes a b-galactosidase reporter. Mice were housed collectively (3 animals per cage) and maintained under a light-dark cycle (12 h light and 12 h dark) with a controlled humidity and temperature, and were feed ad libitum using a standard chow diet. The described experimental procedures were carried out in accordance to the European Community Council Directive 2010/63/EU and approved by the local Ethics Committee for Animal Experiments (Comitè Ètic d’Experimentació Animal-Parc de Recerca Biomèdica de Barcelona, CEEA-PRBB). Mouse genotyping was conducted on tail biopsies as described ^18^ using published primers. In all analysis, WT littermates were used as controls.

### Isolation of mouse pancreatic islets

Mouse pancreata from adult mice (9-14 week old) were perfused with a solution containing 1 mg/mL collagenase-P (Roche) in Hank’s buffered salt solution (HBSS) (Invitrogen) by injection in the common hepatic bile duct. Pancreata were then removed and digested for 12-15 min at 37°C in a water bath. Subsequently, dissociated pancreata were passed through a metal sieve and washed with HBSS. Islets were then isolated using a density gradient purification with HISTOPAQUE-1077 (Sigma), washed and handpicked. Isolated islets were cultured for 24h in RPMI 1640 medium supplemented with 10% FBS, 100 U/mL penicillin, 100 μg/mL streptomycin, 10 mM, HEPES, 1 mM sodium pyruvate and 0.05 mM 2-mercaptoethanol.

### Insulin and glucagon secretion in mouse islets *ex vivo*

24h after isolation, ten to fifteen islets from each mouse in triplicates were used to assess insulin and glucagon secretion. First, islets were pre-incubated for 1 h in Krebs-Ringer bicarbonate (KRB) buffer containing 0.3% bovine serum albumin (BSA) and 2.8 mM glucose. Next, islets were sequentially incubated with the same buffer containing 2.8 mM glucose followed by 20 mM glucose for 1 h. For amino acid-stimulated insulin secretion, islets were pre-incubated for 1 h in KRB buffer containing 0.3% BSA followed by sequential incubations with 0 mM or 12 mM of an amino acid mix containing alanine, arginine, leucine, lysine, proline and glutamic acid. To assess glucagon secretion, islets were pre-incubated for 1 h KRB buffer containing 0.3% BSA and 2.8 mM glucose followed by sequential incubations with 20 mM and 1 mM glucose for 1 h. After these incubations, supernatants were collected to access insulin and glucagon secretion and the remaining islets were sonicated in an alcohol-acid solution to measure total insulin and glucagon content using Mouse Insulin and Glucagon ELISA Kits (Mercodia).

### Measurements of blood glucose and plasma hormones in mice

Adult WT and *Srrm3* mutant (Srrm3 +/−and Srrm3 −/−) mice were fasted during 4h after which blood samples were collected from a tail snip by gentle massaging. After that, mice were offered food and blood sampling was performed 30 min after food intake. Blood glucose levels were measured using a standard glucometer (Glucomen Aero 2K), while circulating insulin and glucagon was measured from plasma using Ultra Sensitive Mouse Insulin and Glucagon ELISA kits (Mercodia).

### Intraperitoneal Glucose and Insulin Tolerance Tests (IPGTT and ITT) in mice

To test glucose tolerance, adult WT and Srrm3 +/−mice were fasted during 12h before they received an intraperitoneal administration of 1g/kg glucose load. Blood glucose levels was measured before (0 min) and 15, 30, 60 and 120 min after glucose injection, from a tail snip by gentle massaging using a blood glucose meter (Glucomen Aero 2K). To test insulin tolerance, mice were restricted to food during 4 h before they received an intraperitoneal administration of 1U/kg insulin (Actrapid, recombinant human insulin, Novo Nordisk), and their blood glucose was measured before (0 min) and 15, 30, 60 and 120 min after insulin administration.

### Immunofluorescence analysis of pancreatic slices

Pancreata from newborn (P1) and 4 weeks old mice were paraffin embedded and cut in 4 μm sections, dewaxed, and incubated with primary antibodies overnight at 4 ºC after microwave antigen retrieval. Primary antibodies used were: Polyclonal Guinea Pig Anti-Insulin (1:200; DAKO, Agilent) and Polyclonal Rabbit anti-Glucagon (1:200; Cell Signaling). IgG secondary antibodies used were: Cy3-conjugated donkey-α-guinea pig and Cy2-conjugated mouse-α-rabbit (1:500; Jackson ImmunoResearch). Slides were imaged by confocal microscopy using a Leica TCS SP5 and images processed by LAS X (Leica) or ImageJ software.

### Analysis of SRRM3 expression and Exon Set Enrichment analyses in human and mouse islets

Transcriptomic analyses in Figs. 7 and Extended Data Fig. 7 aimed at characterizing the expression of *SRRM3* and the relative inclusion of IsletMICs in human or mouse islets comparing two specific conditions (e.g. T2D patients vs controls). Gene expression quantifications were obtained by mapping RNA-seq reads with Salmon^43^ to human or mouse transcriptome (Ensembl release 104). Single cell RNA-seq clustering and cell type classification from all human studies and donors were used to deconvolute human islet bulk RNA-seq with MuSiC^54^. Briefly, raw gene counts from all bulk RNA-seq samples were used as input for MuSiC, as recommended, and samples were filtered based on the proportion of pancreatic islet cell types (Alpha, Beta, Gamma and Delta >60% and only Alpha + Beta >50%; details for each discarded sample is provided in Supplementary Table 1). For *SRRM3* expression, gene raw read counts were normalized with VST from DESeq2. We tested if the *SRRM3* normalized expression differs between groups with a two tailed Wilcoxon test.

To assess the level and direction of global misregulation of the IsletMIC program, we implemented a modification of GSEA, which we termed Exon Set Enrichment Analyses (ESEA). In brief, exon PSI levels were quantified as described above, using *vast-tools* v2.5.1^8^, and ΔPSI for each pairwise comparison (PSI_test - PSI_control) was calculated for those exons with sufficient read coverage in both conditions (score VLOW or higher in least of 60% of all samples per group tested, except for^30^ where events were considered if at least 3 samples per group had coverage greater than VLOW, as provided by *vast-tools*). Next, we generated a background set of alternatively spliced exons for each pairwise comparison, corresponding to exons with sufficient read coverage and a PSI between 5 and 95 in at least one of the conditions, and sorted them based on ΔPSI (rnk file). For each comparison, the tested exon set corresponded to the IsletMICs with sufficient read coverage (gmx file). The ranked backgrounds and the test sets were used as input for *fgsea*^55^.

### Mapping of *SRRM3* regulatory elements and GWAS variants associated with fasting glucose and type 2 diabetes

Position and p-values for GWAS variants associated with “Type 2 diabetes” or “Fasting Glucose” located in the *SRRM3* region were obtained from the databases Finngen (r5.finngen.fi) and CMDKP (hugeamp.org), respectively (date 2021-12-05). Annotated regulatory elements and chromatin marks including pcHi-C and HiChip interactions between clustered enhancers and gene promoter as well as H3K27ac signals under low and high glucose in human islets acting on *SRRM3* were obtained from^4^.

We generated quantile-quantile (Q-Q) plots to examine genomic inflation in IsletMICs or surrounding intronic regions variants for T2D risk and glycemic traits variation. For T2D, we used BMI-adjusted T2D summary statistics^25^. For glycemic traits, we relied on summary statistics data from a trans-ancestral meta-analysis for fasting glucose (FG)^26^. We included genetic variants with MAF< 1% that were intersected with genomic regions comprising 500nt upstream and downstream of IsletMICs.

### Luciferase reporter assay

For episomal reporter assays in INS-1E cells, a 500 nt region of *SRRM3* intron 1 containing the rs67070387 SNP was first amplified from genomic DNA with primers (Supplementary Table 7) containing KpnI/XhoI restriction sites. The amplicon was next cloned into the PGL4.23[luc2/minP] Luciferase Reporter Vector (Promega). Briefly, the amplicon and the vector were simultaneously digested. Next, the vector was dephosphorylated with FastAP (Thermo Scientific). The DNA was then purified and ligated with a T4 DNA ligase (Promega). Next, the generated Reporter Vectors were transformed into E. coli (DH5α) and purified using the Plasmid Maxi Kit (Qiagen). Site-directed mutagenesis was used to introduce single nucleotide variants into the generated construct. The variants were generated by PCR using phosphorylated primers described in Supplementary Table 7. The parental supercoiled double-stranded DNA was digested with *Dpn*I (NEB, 174R0176S) 1h at 37°C and the constructs were transformed in competent E. coli cells (DH5α) by thermal shock. Finally, the introduced variants were validated by Sanger sequencing.

INS-1E cells were transfected in 24 well plates, at a density of 300,000 cells per well, with 200 ng of reporter vectors with or without the insert, plus 20 ng of phRL-CMV Renilla-luciferase to control for transfection efficiency. Transfections were performed with lipofectamine 2000 (Invitrogen) for 8h, according to manufacturer instructions. After 48 h, cells were processed using the Dual Luciferase Assay (Promega) and luciferase activity measured using a VICTOR Multilabel Plate Reader (PerkinElmer), following manufacturer instructions. Firefly luciferase activity was normalized to Renilla luciferase activity.

## Data collection and statistical methods

Insulin secretion data represents samples of independent cell line experiments or mouse islet preparations. Transcriptomics data represents data on the level of single tissues or cells, which are pooled from three independent experiments. Morphological data represents population-wide observations from independent experiments. ATP and actin cytoskeleton measurements represent recordings from individual cells in independent experiments. In vivo data is derived from independent animals. Statistical methods used are represented in each figure legend, statistical tests are two-sided in all cases.

For boxplots: center line, median; box limits, upper and lower quartiles; whiskers, 1.5x interquartile range; points, outliers). Error bars correspond to (SD, SE, CI) unless otherwise stated in the figure caption.

## Data availability

RNA-sequencing data was submitted to GEO under the project GSE198906. All other RNA-seq samples used in this study are publicly available and listed in Supplementary Table 1.

## EXTENDED DATA FIGURES

**Extended Data Fig. 1.**
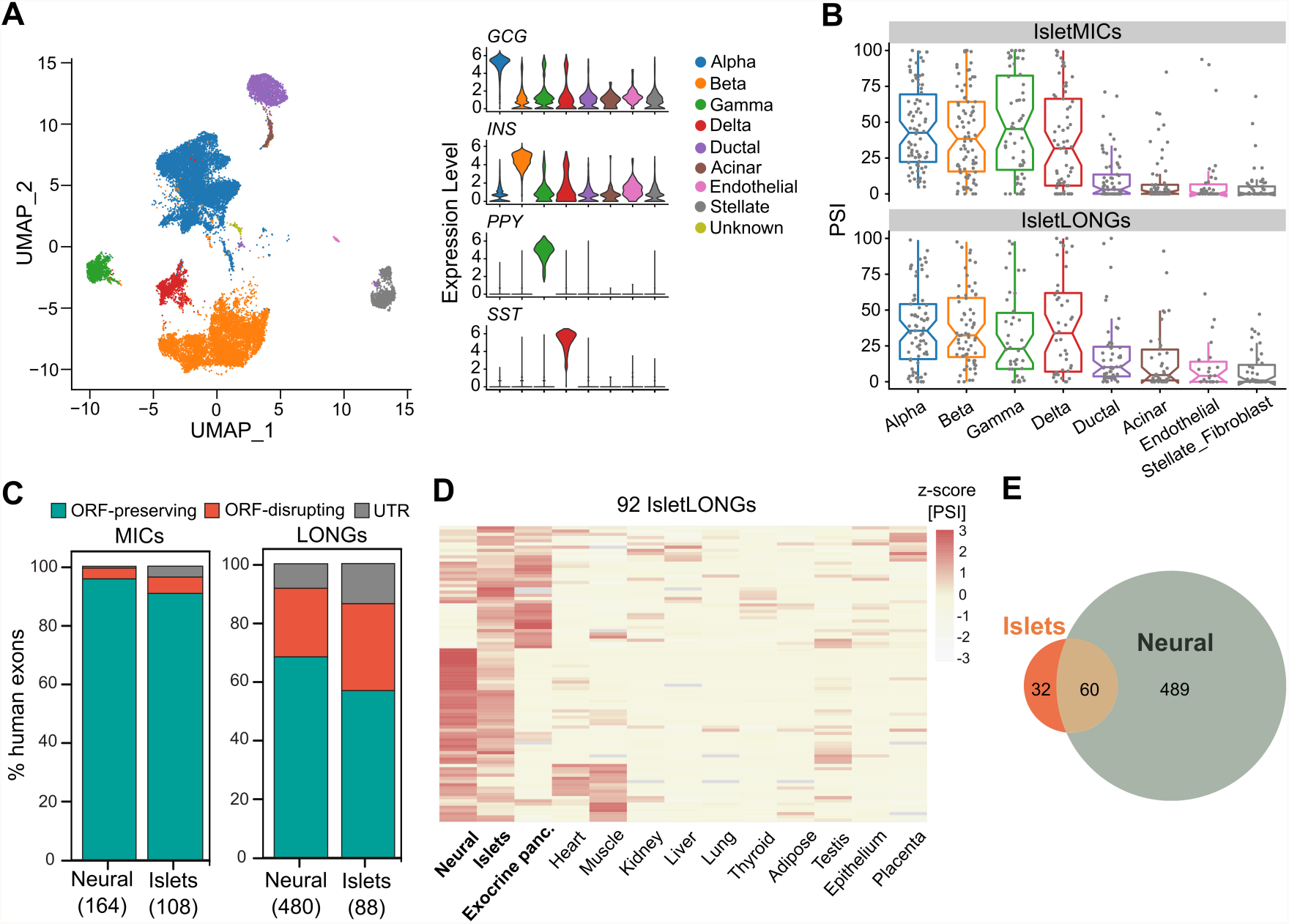
IsletLONGs display a different tissue inclusion pattern and conservation compared to IsletMICs. **A)** UMAP projections of single-cell RNA-seq (scRNA-seq) data from human islets and expression levels of pancreatic hormones according to cell type. **B)** Mean inclusion levels of IsletMICs and IsletLONGs in the different islet cell types from scRNA-seq. **C)** Predicted impact of IsletMICs and IsletLONGs on protein sequences. IsletMICs are largely predicted to generate new protein isoforms upon inclusion by preserving the open reading frame (ORF), while IsletLONGs have a higher proportion of events that disrupt ORFs (i.e. the exon leads to a frame shift and/or introduces a premature termination codon when included or excluded) or impact UTRs. **D)** Heatmap showing the tissue inclusion levels (expressed as z-score of the PSIs) of human cassette exons ≥28 nt enriched in pancreatic islets (IsletLONGs). **E)** Overlap between IsletLONGs and neuron-enriched cassette exons ≥28 nt, showing that 39% (in contrast to 3% of IsletMICs) of tissue-enriched exons are not shared between islet and neural tissues.

**Extended Data Fig. 2.**
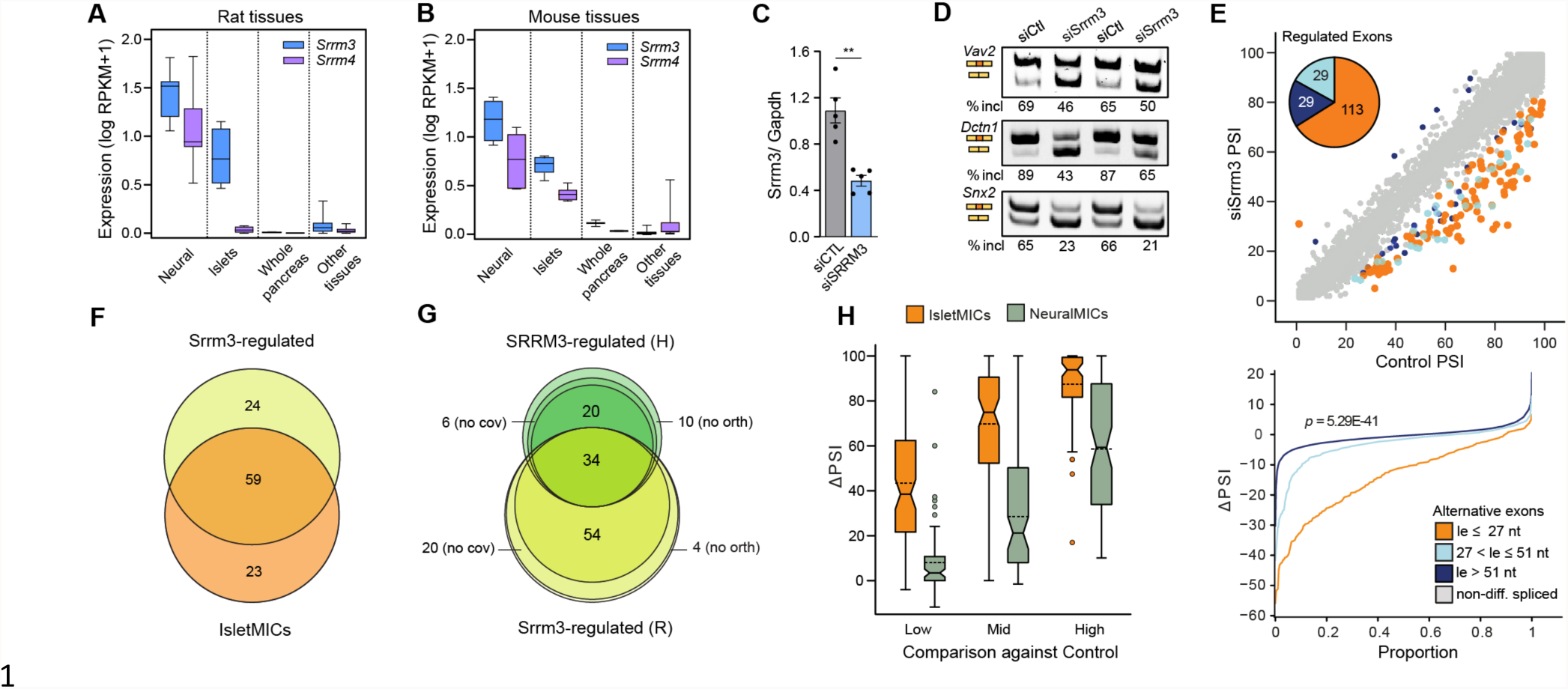
IsletMICs correspond to the subset of neuronal microexons with high sensitivity to *SRRM3*. **A-B)** mRNA expression of the neural microexons regulator *Srrm4* and of its paralog *Srrm3* in neural tissues, pancreatic islets, exocrine pancreas and other rat (A) and mouse (B) tissues (data from VastDB, *n* = 4-20 tissue samples). **C-E)** siRNA-mediated knockdown (KD) of *Srrm3* in INS-1E rat beta cell line. **C)** *Srrm3* mRNA levels measured by qPCR following 48h transfection with *Srrm3* (siSrrm3) or control (siCTL) siRNAs, normalized to Gapdh (*n* = 5, mean ± s.e.m). **, p < 0.01 versus siCTL; paired t test. **D)** RT-PCR assays for selected microexons in control and *Srrm3* KD cells. The positions of inclusion/skipping isoforms and the percentage of microexon inclusion are indicated for two biological replicates. **E)** Global impact of *Srrm3* KD on exon inclusion levels estimated by PSI values from RNA-seq data. Differentially included exons in *Srrm3* KD vs control are shown in orange (microexons, length [le] ≤ 27 nt), light blue (short exons, 27 < le ≤ 51 nt) and dark blue (cassette exons, le > 51 nt). The pie chart shows the number of misregulated exons (|ΔPSI| > 15) according to their size range. Lower panel shows ΔPSI cumulative proportions for microexons, short exons and alternative cassette exons. P-value was obtained from two-sided Wilcoxon test comparing the distributions of microexons and cassette exons of length > 51 nt. PSI values are the mean of three independent experiments. **F)** Overlap between *Srrm3*-regulated microexons in rat INS-1E cells and rat IsletMICs. Only microexons with sufficient read coverage in both comparisons are shown. **G**) Overlap between *SRRM3*-regulated microexons in rat INS-1E (R) and human EndoC-βH1 (H). Microexons presenting no ortholog (“no orth”) or with no sufficient read coverage (“no cov”) in the other species are indicated. **H)** *SRRM3* overexpression in HeLa cells at three different levels reveals different sensitivities for IsletMICs and neuronal-only MICs (NeuralMICs). Exon inclusion leveIs were quantified from RNA-seq and differences in inclusion between control and *SRRM3*-overexpressing cells are shown for IsletMICs and NeuralMICs (*n* = 1).

**Extended Data Fig. 3.**
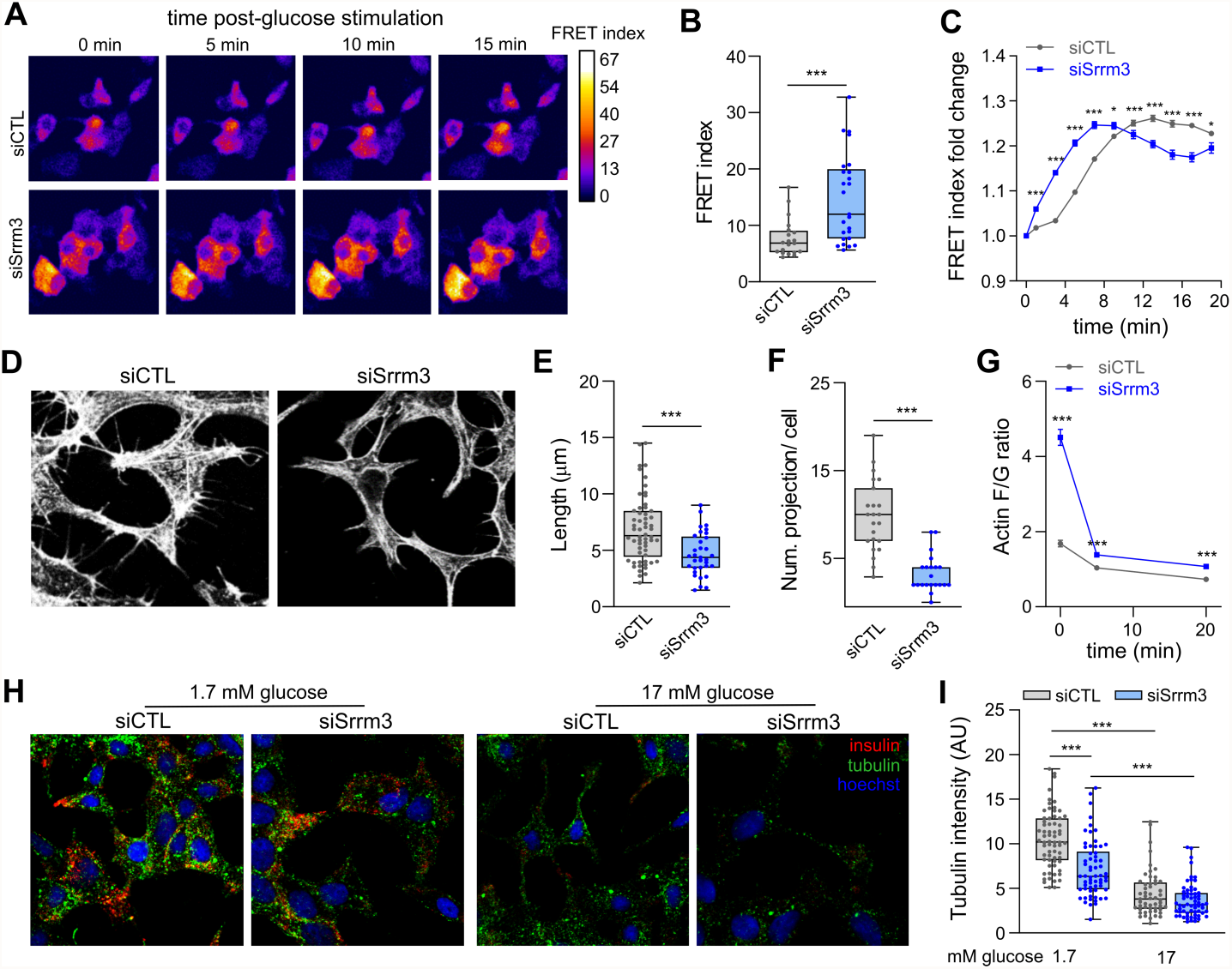
*Srrm3* regulates insulin secretory functions of pancreatic beta cells. **A)** Representative images of control and *Srrm3*-depleted cells FRET measurements of the levels of ATP produced at 1.7 mM glucose (0 min) and following stimulation with 17 mM glucose (5-15 min). **B)** FRET measurements of ATP production at 1.7 mM glucose indicating that *Srrm3*-depleted cells display increased basal cellular metabolism (*n* = 20 individual cells). **C)** Fold increase in ATP production following exposure to high glucose (17 mM) showing that *Srrm3* KD increases ATP production dynamics (*n* = 20 individual cells). **D)** Representative confocal images of control and *Srrm3* KD cells actin filaments. **E, F)** Quantification of the length (E) and number of projections per cell (F) of actin-rich filopodia (*n* = 20 individual cells). **G)** Quantification of glucose-induced actin depolymerization measured by the intensity ratio between filamentous [F] and globular [G] actin (*n* = 94 individual cells). F and G actin were stained with Atto 550-phalloidin and Alexa 488-DNase I, respectively. **H)** Representative confocal images of control and *Srrm3* KD cells stained against β-tubulin (green) and insulin (red) following 1h exposure to 1.7 mM and 17 mM glucose. **I)** Fluorescence intensity of immunostained tubulin in low and high glucose concentrations showing that *Srrm3* KD cells present a lower microtubule nucleation at low glucose compared to control cells (n =53-66 individual cells). All statistical comparisons: *, p < 0.05; **, p < 0.01 and ***, p < 0.001 versus siCTL or CTL ASO; unpaired t test.

**Extended Data Fig. 4.**
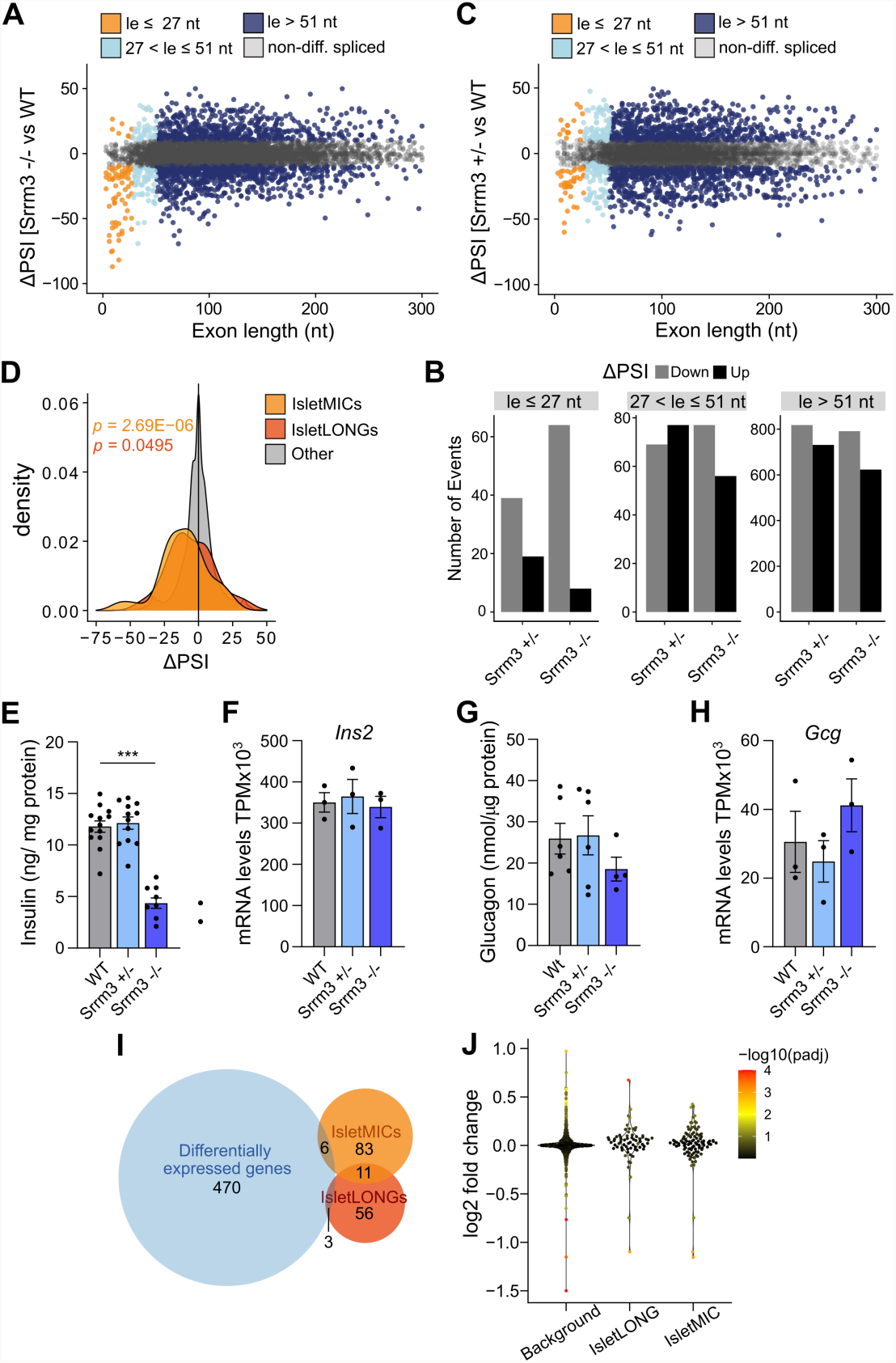
*Srrm3* depletion in mouse islets causes IsletMIC downregulation and increased stimulated insulin release. **A)** Differences in inclusion levels (ΔPSI) in islets from *Srrm3* −/− vs wild type (WT) mice for all exons shorter than 300 nucleotides with sufficient read coverage. PSI values are the mean of three mice islet preparations. **B)** Number of misregulated alternative exons according to size range in *Srrm3* −/− and *Srrm3* +/− mouse islets compared to WT ones. Black and grey bars indicate exons with ΔPSI > 15 and ΔPSI < −15, respectively. **C)** Differences in inclusion levels (ΔPSI) in islets from Srrm3 +/− vs wild type (WT) mice for all exons shorter than 300 nucleotides with sufficient read coverage. PSI values are the mean of three mice islet preparations. **D)** Density plots for ΔPSI distributions in *Srrm3* +/ islets of IsletMICs, IsletLONGs and other alternative exons. P-values were obtained from two-sided Wilcoxon test comparing the distributions of IsletMICs (orange) or IsletLONGs (red) against other alternative exons. **E)** Total insulin protein content in mice islets measured by ELISA (*n* = 10-13, mean ± s.e.m). **F)** Insulin mRNA levels in mice islets measured by RNA-seq (*n* = 3, mean ± s.e.m). **G)** Total glucagon protein content in mice islets measured by ELISA (*n* = 4-6, mean ± s.e.m). **H)** Glucagon mRNA levels in mice islets measured by RNA-seq (*n* = 3, mean ± s.e.m). Statistical comparisons in (E-H): ***, p < 0.001 versus WT; unpaired t test. **I)** Overlap between differentially expressed genes in *Srrm3* −/− islets and genes harbouring IsletMIC and IsletLONG alternative exons. **J)** Distribution of log2 fold change values in genes containing either IsletMICs or IsletLONGs compared to background (genes expressed in mouse islets). Color code represents −log10 adjusted p values.

**Extended Data Fig. 5.**
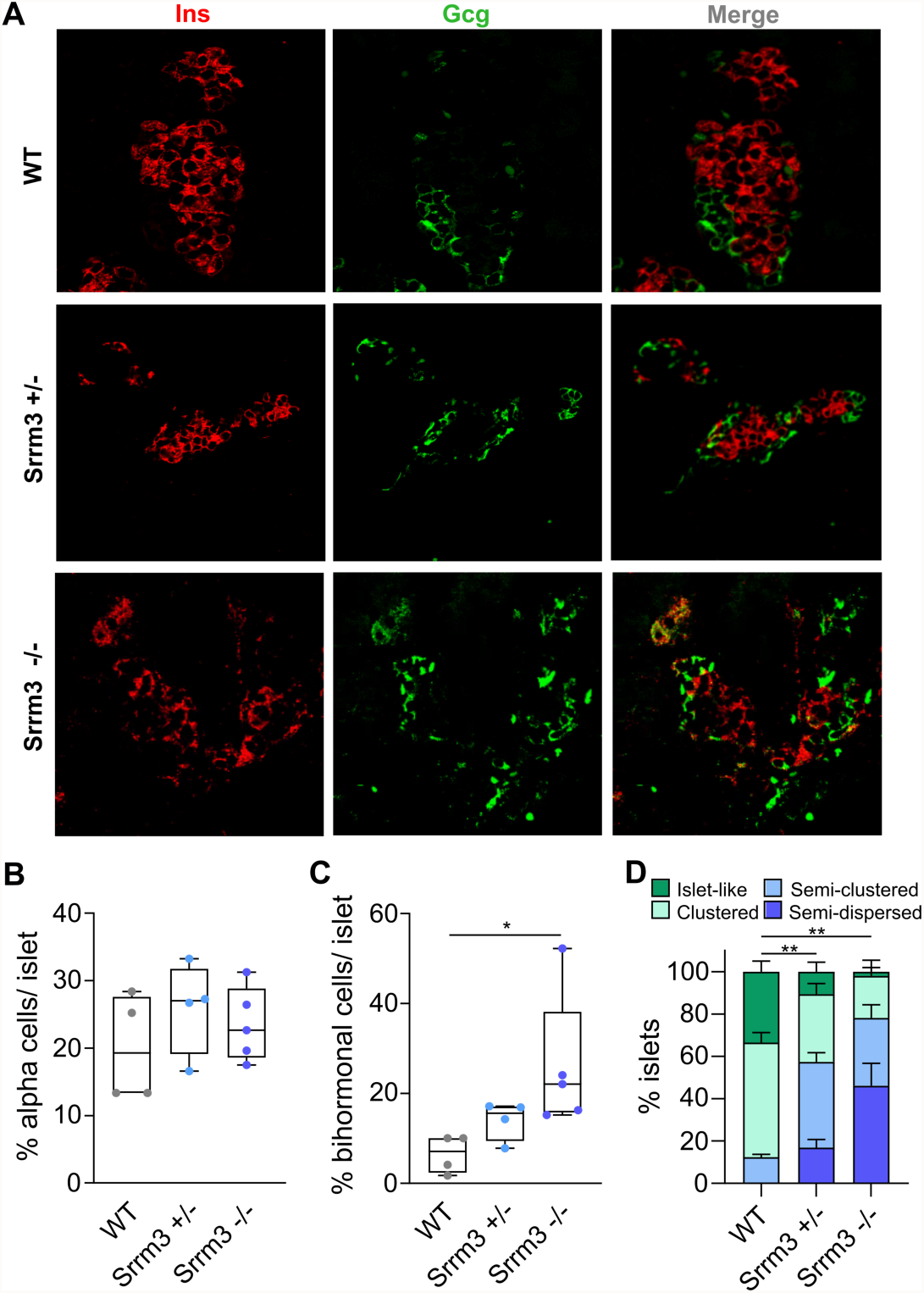
*Srrm3* depletion disrupts islet cell composition and architecture. **A)** Representative immunofluorescence images of islets from WT and *Srrm3* mutant neonatal mice stained for insulin (Ins; red) and glucagon (Gcg; green). **B**,**C)** Quantification of the fraction of alpha cells (B) and insulin and glucagon double-positive cells (C) (*n* = average of 9-12 islets from 4-5 mice per genotype including males and females). **D)** Percent of endocrine cell clusters in well-organized islet-like structures or displaying more dispersed organization. Statistical comparisons: *, p < 0.05 and **, p < 0.01 versus WT; unpaired t test.

**Extended Data Fig. 6.**
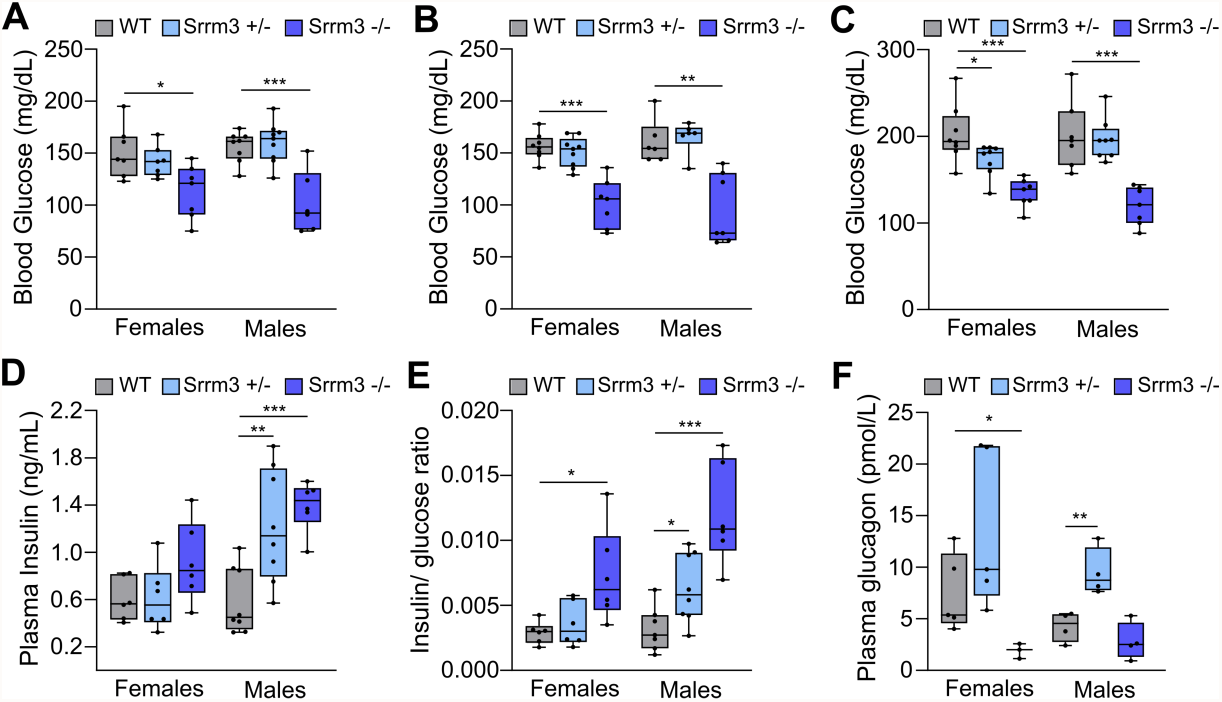
*Srrm3* mutant mice display hypoglycemic hyperinsulinemia. Glycemia and plasma insulin and glucagon levels segregated by sex. **A)** Random blood glucose measurements in *Srrm3* mutant and wild type mice fed ad libitum (*n* = 7-9). **B)** Blood glucose levels following a 4h fast (*n* = 7-9). **C)** Blood glucose levels at 30 min postprandial (*n* = 7-9). **D**,**E)** Insulin plasma levels (D) and ratio between plasma insulin and blood glucose (E) at 30 min postprandial (*n* = 6-8). **F)** Plasma glucagon at 4h fasting (*n* = 4-5). Statistical comparisons: *, p < 0.05; **, p < 0.01 and ***, p < 0.001 versus WT; unpaired t test.

**Extended Data Fig. 7.**
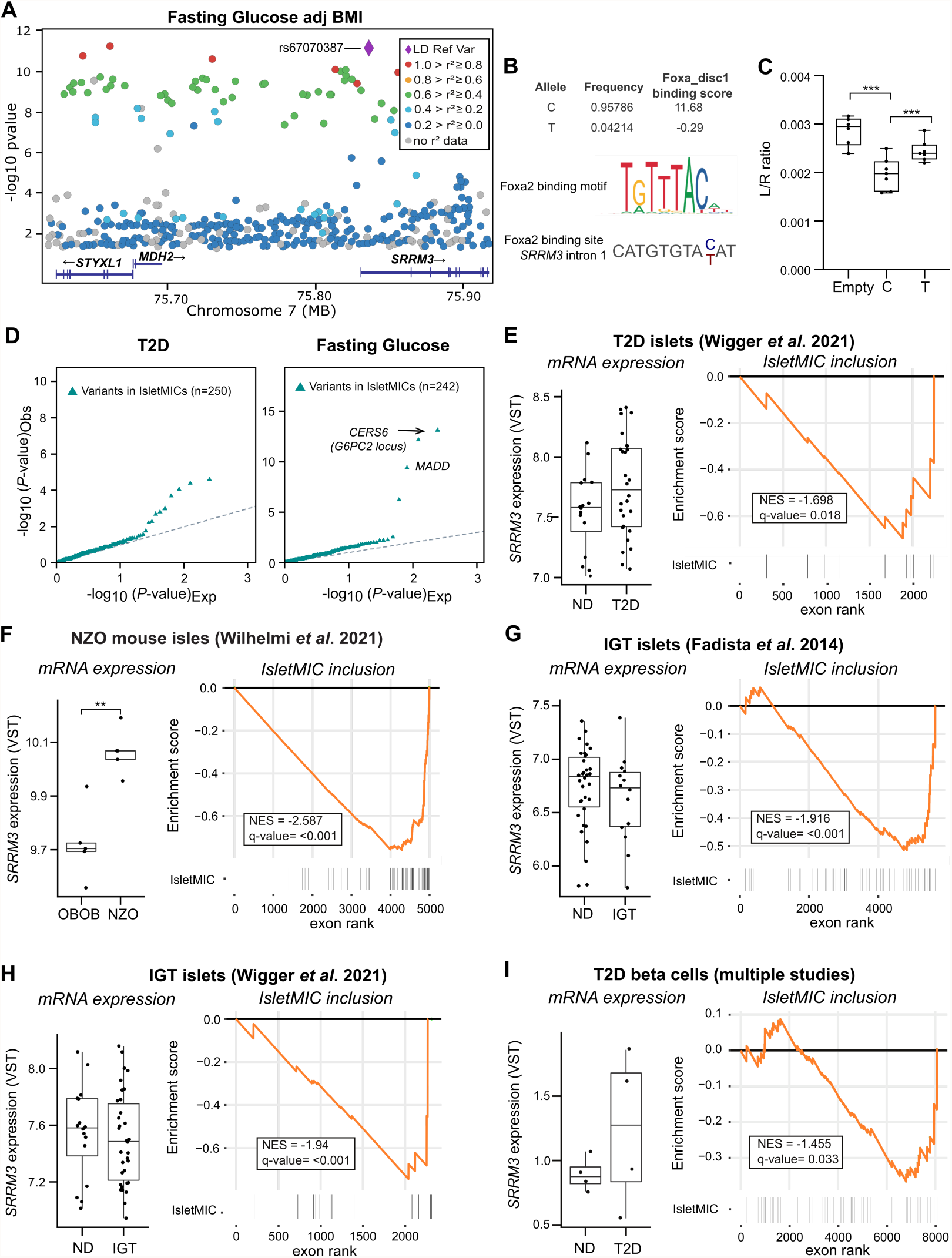
The *SRRM3* locus responds to glucose and harbors genetic variants associated with fasting glucose and type 2 diabetes risk. **A)** Regional plot of Fasting Glucose GWAS variants from the CMDKP database (hugeamp.org). **B)** Allele frequency and predicted impact on a Foxa2 binding motif for the SNP rs67070387 associated with elevated Fasting glucose. **C)** Luciferase assay in INS-1E cells following transfection of a control reporter vector or carrying the genomic sequence surrounding rs67070387 with one or the other allele (*n* = 6-7). ***, p < 0.001 versus indicated by unpaired t test. **D)** Quantile-quantile (QQ) plots showing distribution of p-values associated with type 2 diabetes from^25^ or fasting glucose from^26^ for variants located in 1Kb genomic regions containing microexons. **E)** *SRRM3* expression and IsletMIC inclusion in human islets from non-diabetic (ND) and type 2 diabetes (T2D) individuals from^30^ (*n* = 16-28). **F)** *Srrm3* expression and IsletMIC inclusion in islets from B6-ob/ob (OBOB) and New Zealand Obese (NZO) mice from^31^ (*n* = 5). **G)** *SRRM3* expression and IsletMIC inclusion in human islets from non-diabetic (ND) and impaired glucose tolerant (IGT) individuals from^28^ (*n* = 14-34). **H)** *SRRM3* expression and IsletMIC inclusion in human islets from non-diabetic (ND) impaired glucose tolerant (IGT) individuals from^30^ (n = 16-34). **I)** *SRRM3* expression and IsletMIC inclusion in single beta cells from non-diabetic (ND) and type 2 diabetes (T2D) individuals from multiple studies (*n* = 4) (see methods).

